# Origin and microenvironment contribute to the sexually dimorphic phenotype and function of peritoneal macrophages

**DOI:** 10.1101/837336

**Authors:** Calum C. Bain, Douglas A. Gibson, Nicholas Steers, Katarina Boufea, Pieter A. Louwe, Catherine Docherty, Victor Huici, Rebecca Gentek, Marlene Magalhaes-Pinto, Marc Bajenoff, Cecile Benezech, David Dockrell, Philippa TK Saunders, Nizar Batada, Stephen J Jenkins

## Abstract

Macrophages reside in the body cavities where they maintain serosal homeostasis and provide immune surveillance. Peritoneal macrophages are implicated in the aetiology of pathologies including peritonitis, endometriosis and metastatic cancer thus understanding the factors that govern their behaviour is vital. Using a combination of fate mapping techniques, we have investigated the impact of sex and age on murine peritoneal macrophage differentiation, turnover and function. We demonstrate that the sexually dimorphic replenishment of peritoneal macrophages from the bone marrow, which is high in males and very low in females, is driven by changes in the local microenvironment that arise upon sexual maturation. Population and single cell RNAseq revealed striking dimorphisms in gene expression between male and female peritoneal macrophages that was in part explained by differences in composition of these populations. By estimating the time of residency of different subsets within the cavity and assessing development of dimorphisms with age and in monocytopenic *Ccr2*^−/−^ mice, we demonstrate that key sex-dependent features of peritoneal macrophages are a function of the differential rate of replenishment from the bone marrow while others are reliant on local microenvironment signals. Importantly, we demonstrate that the dimorphic turnover of peritoneal macrophages contributes to differences in the ability to protect against pneumococcal peritonitis between the sexes. These data highlight the importance of considering both sex and age in susceptibility to inflammatory and infectious disease.

## Introduction

Macrophages are present in every tissue of the body, where they provide immune protection and orchestrate tissue repair following insult or injury. Peritoneal macrophages are arguably the most studied population of macrophages in the body, having been used extensively as a convenient source of macrophages for *ex vivo* analyses for decades. Despite this, the heterogeneity of peritoneal macrophages and much of the biology that governs their development, differentiation and function remains unclear. Macrophages in the peritoneal cavity are programmed for ‘silent’ clearance of apoptotic cells, maintenance of innate B1 cells through secretion of CXCL13, and for immune surveillance of the cavity and neighbouring viscera ^1–4^. However, they are also implicated in many pathologies, including peritonitis, endometriosis, post-surgical adhesions, pancreatitis and metastatic cancer ^5–15^, although the exact role(s) they play in these processes is not fully understood.

Under physiological conditions, at least two macrophage populations are present in the murine peritoneal cavity, with those expressing high levels of F4/80, CD11b and CD102 outnumbering their F4/80^lo^MHCII^+^ counterparts by approximately 10-fold. F4/80^hi^CD102^+^ macrophages (sometimes referred to as ‘large’ peritoneal macrophages ^16^) rely on the transcription factors C/EBPβ and GATA6 for their differentiation and survival ^17–20^, with the latter under the control of retinoic acid proposed to derive, in part, from the omentum ^19^. In contrast, F4/80^lo^MHCII^+^ macrophages (sometimes referred to as ‘small’ peritoneal macrophages) rely on IRF4 for their differentiation and can be further defined by their expression of CD226 and the immunomodulatory molecule RELMα ^21, 22^. Notably, recent studies employing lineage tracing techniques have established that F4/80^lo^MHCII^+^ macrophages arise postnatally, are short-lived and replaced by Ly6C^hi^ classical monocytes in a CCR2-dependent manner ^20–23^. In contrast, F4/80^hi^CD102^+^ macrophages are longer-lived cells that originally derive from embryonic sources, but are subsequently replaced by cells of haematopoietic stem cell (HSC) origin ^21, 24^. Importantly, we have recently shown that unlike resident macrophages in numerous other tissues the turnover of peritoneal F4/80^hi^CD102^+^ macrophages from the bone marrow is highly sex-dependent, with high and low rates in male and female mice respectively ^21^. We have also shown that long-lived macrophages can be identified by their expression of the phagocytic receptor, Tim4, whereas most recent descendants of BM-derived cells amongst the F4/80^hi^CD102^+^ macrophage compartment are Tim4^−21^. Indeed Tim4 expression has been shown to a feature of long-lived macrophages in other tissues ^25–29^. However, it remains unclear if further heterogeneity exists amongst these broadly-defined populations and if sexually-dimorphic turnover influences the composition and function of the F4/80^hi^CD102^+^ macrophage population in other ways.

Sex is a variable often overlooked in immunological research ^30^ despite strong sex biases in many pathologies including autoimmune disorders and infection susceptibility ^31^. Notably, sex dimorphisms in the immune system are present a diverse range of species from insects, bird, lizards and mammals ^31^, demonstrating this is an evolutionary conserved phenomenon. It is therefore essential to understand how intrinsic factors such as sex control the behaviour of innate immune effector cells. Specifically, sex has been proposed to affect macrophage behaviour, such as influencing the differentiation of brain microglia ^32–34^ and sex hormones appear able to directly regulate gene expression ^35^ and proliferation ^36^ of macrophages. While previous studies have considered the effects of sex on peritoneal macrophage behaviour, many of these have focussed on *in vitro* functional assessments using macrophages elicited by injection of an irritant or inflammatory agent ^37^, or have not appreciated the complexity of the peritoneal macrophage compartment ^36, 38^.

Here we have used a combination of fate-mapping techniques together with population-level and single cell RNA sequencing (scRNAseq) to dissect the role of sex in the composition, environmental imprinting and function of peritoneal macrophages. We show that the F4/80^hi^CD102^+^ macrophage population is heterogeneous and that dimorphic turnover is associated with divergence in the heterogeneity of this compartment with age. Specifically, we demonstrate that the sexual dimorphism in replenishment from the bone marrow and phenotype arise following sexual maturation. Furthermore, we provide examples of transcriptional and functional dimorphisms that arise due to sex differences in turnover versus those arising directly from sex differences in the peritoneal microenvironment. Importantly, we identify the C-type lectin receptor CD209b (also known as Specific ICAM3-grabbing nonintegrin-related 1; SIGN-R1) as a marker whose expression is determined by replenishment that becomes increasingly dimorphic with age, and show that sex-dependent resistance to pneumococcal peritonitis arises, in part, due to dimorphic expression of CD209b.

## Results

### Environment drives sexual dimorphism in macrophage replenishment in the peritoneal cavity

We first set out to determine if the dimorphic effects in peritoneal macrophage replenishment were due to the peritoneal environment or to cell-intrinsic differences in the ability of male and female monocytes to generate F4/80^hi^CD102^+^ macrophages at this site. To this end, we generated sex-mismatched, tissue-protected bone marrow (BM) chimeric mice to measure the turnover of peritoneal F4/80^hi^CD102^+^ macrophages from the BM (Figure 1a) and assess the role of sex in this process. Wild type (CD45.1/.2^+^) mice were irradiated, with all but the head and upper torso protected with lead to prevent direct exposure to ionising radiation (Figure 1b), before being reconstituted with sex-matched (female > female, male > male) or sex-mismatched (female > male, male > female) BM. Following at least 8 weeks reconstitution, the non-host chimerism was measured in peritoneal macrophages. Consistent with our previous work ^21^, only low levels of non-host chimerism could be detected amongst peritoneal F4/80^hi^CD102^+^ macrophages from female > female BM chimeric mice, whereas high levels were detected in their male > male counterparts (Figure 1c&d), confirming marked sex dimorphism in macrophage turnover. Importantly, this dimorphism was specific to F4/80^hi^CD102^+^ peritoneal macrophages, as all other leukocyte subsets showed identical replenishment in male and female BM chimeric mice (**Supplementary Figure 1a**). Strikingly, F4/80^hi^CD102^+^ peritoneal macrophages from sex mismatched (female > male) chimeras had similar levels of chimerism to male > male chimeras (Figure 1c&d), demonstrating that female and male monocytes have equal ability to generate F4/80^hi^CD102^+^ macrophages in the male peritoneal cavity. Female recipients rejected male BM and thus chimerism in this group could not be determined.

**Figure 1.**
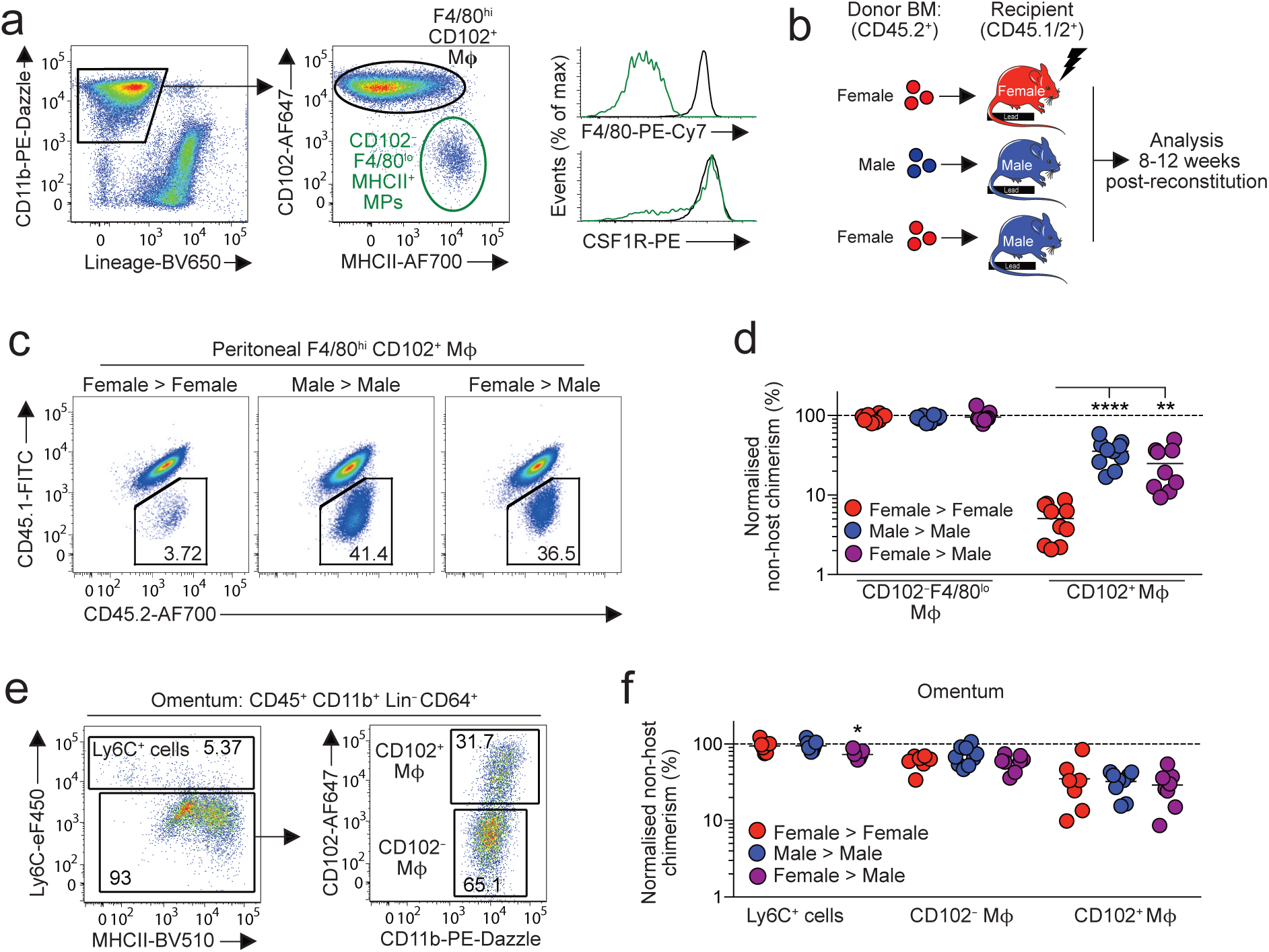
Environment drives sexual dimorphism in macrophage replenishment in the peritoneal cavity. (**a**) Expression of CD11b and CD3, CD19, Ly6G and SiglecF (‘Lineage’) by live CD45^+^ peritoneal leukocytes (*left*) and expression of CD102 and MHCII by CD11b^+^ Lin^−^ cells (*centre*) from adult C57Bl/6 female mice. Histograms show expression of F4/80 and CSF1R by CD102^+^ and CD102^−^ MHCII^+^ cells. (**b**) Experimental scheme for the generation of sex mis-matched, tissue-protected bone marrow (BM) chimeric mice. (**c**) Representative expression of CD45.1 and CD45.2 by peritoneal F4/80^hi^CD102^+^ macrophages from sex matched or mismatched tissue protected BM chimeric mice 8-12 weeks post-reconstitution. (**d**) Normalized non-host chimerism of peritoneal F4/80^hi^CD102^+^ macrophages from sex matched or mismatched tissue-protected BM chimeric mice 8-12 weeks post-reconstitution. Data are normalised to the non-host chimerism of Ly6C^hi^ monocytes. **P<0.01, ****P<0.0001 One-way ANOVA. (**e**) Gating strategy to identify macrophages amongst omental isolates. Expression of Ly6C and MHCII by CD11b^+^ Lin^−^CD64^+^ cells and expression of CD102 by Ly6C^−^ cells to identify CD102^+^ and CD102^−^ macrophages. (**f**) Normalized non-host chimerism of omental Ly6C^+^ monocytes and CD102^+^ and CD102^−^ macrophages from mice in (**d**). Data are normalised to the non-host chimerism of Ly6C^hi^ monocytes. *P<0.05. One-way ANOVA. Symbols represent individual animals and horizontal bars represent the mean. Data represent 9 (female > male) or 10 (sex matched) mice per group pooled from two independent experiments.

The omentum has been implicated in the differentiation of F4/80^hi^ macrophages in the peritoneal cavity, potentially acting as site of macrophage maturation ^19, 39, 40^. Indeed, CD102^+^ macrophages that co-express GATA6, can be detected amongst omental isolates ^19^, together with CD102^−^MHCII^+^ macrophages and a population of Ly6C^+^ CD11b^+^ cells similar to monocytes (Figure 1e and **Supplementary Figure 1b-c**). To determine if the dimorphic replenishment of peritoneal F4/80^hi^CD102^+^ macrophages arises in the omentum, we assessed non-host chimerism in the macrophage populations within this site. While this showed clear differences in the turnover of CD102-defined macrophage populations from BM, with higher replenishment in the CD102^−^ fraction, no sex dimorphism was detected in any monocyte/macrophage population within the omentum (Figure 1f). Furthermore, the chimerism of omental and peritoneal CD102^+^ macrophages in male recipients was identical, rather than showing the gradation that would have been expected if omental macrophages were intermediate precursors between monocytes and cavity CD102^+^ cells (**Supplementary Figure 1d**). Thus, the sexual dimorphism in peritoneal F4/80^hi^CD102^+^ macrophage replenishment is driven by factors present in the local environment.

### Sexual dimorphism in peritoneal macrophage replenishment occurs following sexual maturity

To extend these findings and to assess macrophage turnover at different stages of maturity, we next used a genetic fate mapping approach. Adoptive transfer experiments suggest F4/80^lo^MHCII^+^ macrophages in the peritoneal macrophage compartment act, in part, as precursors of F4/80^hi^CD102^+^ macrophages ^20^ and we have recently shown that this differentiation can be mapped by exploiting their expression of CD11c ^21^. Thus, in CD11c^Cre^.*Rosa26*^LSL-eYFP^ mice (Figure 2a), in whom active or historic expression of CD11c leads to irreversible labelling with eYFP, labelled cells accumulate with age in the F4/80^hi^CD102^+^ macrophage compartment, despite these cells themselves not actively expressing CD11c ^21^. We therefore used CD11c^Cre^.*Rosa26*^LSL-eYFP^ mice to compare the rate of eYFP^+^ cell accumulation in peritoneal F4/80^hi^CD102^+^ macrophages from male and female mice. In juvenile/prepubescent mice (4 weeks of age), the extent of eYFP labelling was relatively similar between male and female peritoneal F4/80^hi^CD102^+^ macrophages and indeed was marginally higher in female mice (Figure 2b). By 16 weeks of age, the frequency of eYFP^+^ cells amongst F4/80^hi^CD102^+^ macrophages had increased in both male and female mice compared with their 4-week-old counterparts. However, although there was no difference in CD11c protein expression by male and female F4/80^hi^CD102^+^ macrophages (Figure 2c), significantly higher levels of eYFP labelling were detected amongst male peritoneal macrophages, consistent with more rapid accumulation of newly differentiated macrophages in male mice (Figure 2b). Consistent with our previous findings made using tissue-protected BM chimeras^21^, the sexual dimorphism in eYFP labelling was not detected in F4/80^hi^CD102^+^ macrophages from the pleural cavity (Figure 2d), where both male and female pleural cells exhibited high levels of labelling that were equivalent to those seen in the male peritoneal cavity by 16 weeks. Collectively, these data confirm that the peritoneal environment controls macrophage turnover and suggest that dimorphisms arise in this site following sexual maturity.

**Figure 2.**
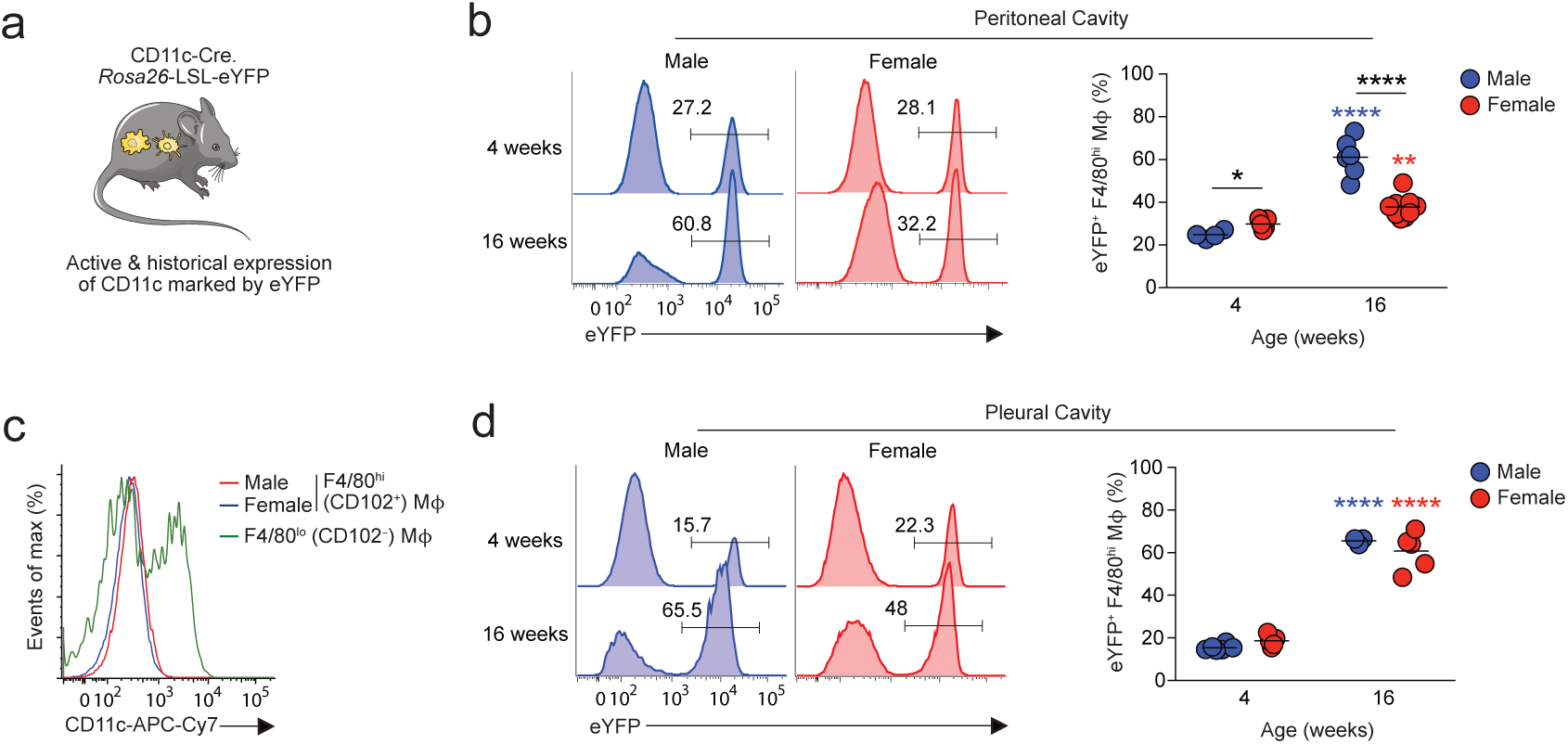
Sexual dimorphism in peritoneal macrophage replenishment occurs following sexual maturity. (**a**) Experimental scheme of CD11c^Cre^.*Rosa26*^LSL-eYFP^ fate-mapping mice. (**b**) Representative expression of eYFP by peritoneal F4/80^hi^CD102^+^ macrophages from male and female CD11c^Cre^.*Rosa26*^LSL-eYFP^ fate-mapping mice at 4 and 16 weeks of age. Right, frequency of eYFP^+^ cells amongst F4/80^hi^CD102^+^ macrophages in male and female mice at the indicated ages. Symbols represent individual animals and horizontal bars represent the mean. Data represent 4 (male 4 weeks), 5 (female 4 weeks), 6 (male 16 weeks) or 9 (female 16 weeks) mice per group pooled from two independent experiments. (**c**) Expression of CD11c by peritoneal F4/80^hi^CD102^+^ macrophages from male and female and CD102^−^ MHCII^+^ cells from female mice. (**d**) Representative expression of eYFP by pleural F4/80^hi^CD102^+^ macrophages from male and female CD11c^Cre^.*Rosa26*^LSL-eYFP^ fate-mapping mice at 4 and 16 weeks of age. Right, frequency of eYFP^+^ cells amongst pleural F4/80^hi^CD102^+^ macrophages in male and female mice at the indicated ages. Symbols represent individual animals and horizontal bars represent the mean. Data represent 3 (male 16 weeks), 5 (female 4 & 16 weeks) or 6 (male 4 weeks) per group pooled from two independent experiments.

### Ovariectomy leads to increased macrophage replenishment

The onset of sexually dimorphic turnover of peritoneal macrophages following sexual maturation and the uniquely slow replenishment of female peritoneal macrophages suggested that factors involved in female reproductive function may drive this dimorphism. Therefore, we next assessed macrophage turnover in females after ovariectomy (OVX). Thus, female > female tissue protected BM chimeric were generated and after 8 weeks reconstitution, both ovaries were surgically removed (bilateral OVX), before measuring non-host chimerism after another 8 weeks. To account for the potential effects of surgery on macrophage replenishment, BM chimeric mice receiving sham surgery or unilateral OVX were used as controls, together with unmanipulated BM chimeric mice. As expected, the cessation of ovarian estradiol production caused by bilateral OVX led to complete atrophy of the uterine horn; this was not seen in mice with unilateral OVX, or in other control groups (Figure 3a). Bilateral OVX had no effect on the numbers of F4/80^hi^CD102^+^ and CD102^−^MHCII^+^ macrophages in the peritoneal cavity when compared with the control groups (Figure 3b). Although bilateral OVX led to increased proportions and absolute numbers of eosinophils, these differences did not attain statistical significance and the opposite pattern was found with B1 cells (**Supplementary Figure 2**). Importantly and in striking contrast to the very low levels of chimerism (∼1%) detected in unmanipulated control chimeras (Figure 3c), sham surgery and unilateral OVX led to significant increases the level of chimerism compared with unmanipulated chimeric mice, demonstrating that minimally-invasive laparotomy itself appears to have long term effects on the dynamics of peritoneal macrophages in female mice. Nevertheless, complete removal of the ovaries further elevated macrophage turnover, with chimerism reaching approximately 12%. No difference was found between the chimerism seen after sham surgery with or without unilateral OVX, indicating that the OVX procedure itself does not exaggerate the effects of laparotomy and that it is the compete loss of ovarian function that underlies the further elevation in macrophage turnover that results from bilateral OVX. Consistent with these results, significantly more Tim4^−^CD102^+^ macrophages were present in the cavity of mice that received bilateral OVX than any other group (Figure 3d), further supporting the idea of elevated macrophage replenishment from BM. Notably, no differences in chimerism or in Tim4-defined subsets could be detected amongst F4/80^hi^CD102^+^ macrophages from the pleural cavity, again confirming that the effect of surgery and ovariectomy on macrophage turnover are specific to the peritoneal cavity (Figure 3b, c).

**Figure 3.**
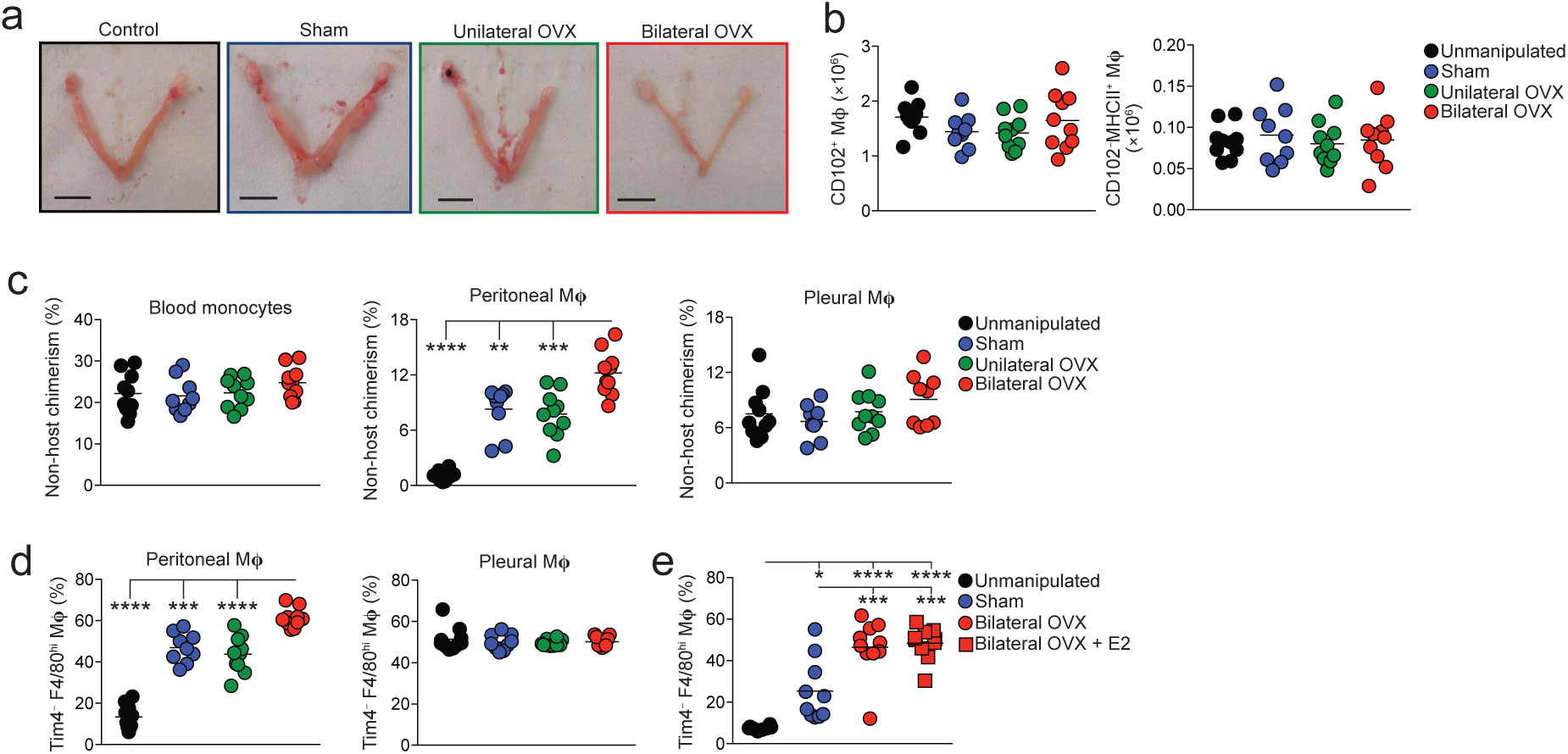
Ovariectomy leads to increased macrophage replenishment. (**a**) Representative images of the uterine horns of tissue-protected BM chimeric mice that had received unilateral or bilateral oophorectomy (OVX), sham surgery or were completely unmanipulated (control). (**b**) Absolute number of F4/80^hi^CD102^+^ macrophages and CD102^−^MHCII^+^ cells obtained from the peritoneal cavity of tissue-protected BM chimeric mice that had received surgery 8 weeks earlier. Symbols represent individual animals and horizontal bars represent the mean. Data represent 9 (sham) or 10 (control, unilateral, bilateral) mice per group pooled from two independent experiments.| (**c**) Non-host chimerism of blood Ly6C^hi^ blood monocytes (*left*) and F4/80^hi^CD102^+^ macrophages obtained from the peritoneal (*centre*) or pleural (*right*) cavity of tissue-protected BM chimeric mice that had received surgery 8 weeks earlier. Symbols represent individual animals and horizontal bars represent the mean. Data represent 9 (sham) or 10 (control, unilateral, bilateral) mice per group pooled from two independent experiments. **P<0.01, ***P<0.001, ****P<0.0001. One-way ANOVA with Tukey’s multiple comparisons test. (**d**) Frequency of Tim4^−^ cells amongst F4/80^hi^CD102^+^ macrophages obtained from the peritoneal (*centre*) or pleural (*right*) cavity of mice in (**b**). ***P<0.001, ****P<0.0001. One-way ANOVA with Tukey’s multiple comparisons test. (**e**) Frequency of Tim4^−^ cells amongst F4/80^hi^CD102^+^ macrophages obtained from the peritoneal cavity of unmanipulated C57Bl/6 female mice (controls) or age-matched females that received bilateral OVX or sham surgery 4 weeks earlier. One group received exogeneous estradiol (E2) thrice weekly for 3 weeks. Symbols represent individual animals and horizontal bars represent the mean. Data represent 8 (control) or 10 (sham, bilateral, bilateral + E2) mice per group pooled from two independent experiments. *P<0.05, **P<0.01, ***P<0.001, ****P<0.0001. One-way ANOVA with Tukey’s multiple comparisons test.

Estrogens are the prototypical female sex steroid hormones which are ablated by OVX. To assess if estrogen influences macrophage replacement, we repeated the OVX experiment with an additional group of bilateral OVX mice receiving exogenous estradiol (E2). However, while this treatment reversed OVX-mediated atrophy of the uterine horns and peritoneal eosinophilia, it had no effect on the heightened rate of replenishment of F4/80^hi^CD102^+^ peritoneal macrophages in OVX mice, suggesting that estradiol is not directly responsible for generating the sex dimorphism in peritoneal macrophage turnover (Figure 3e). As males exhibit much greater levels of adipose tissue in the peritoneal cavity (**Supplementary Figure 3**) and a common feature of ovariectomy/oophorectomy in mice and humans is increased adiposity ^41, 42^, something we also noted in our experiments (data not shown), we combined a high fat diet (HFD) with our BM chimeric system to reveal if changes in adiposity affect replenishment of peritoneal F4/80^hi^CD102^+^ macrophage. However, replenishment was not affected by diet in either males or females, despite the expected increase in body weight and adipose tissue seen in mice on an HFD (**Supplementary Figure 3**). Hence, sexual maturation controls dimorphic turnover in the peritoneal cavity through a mechanism controlled at least in part by the female reproductive system, but independently of estrogen and local fat deposition.

### Sex determines the transcriptional signature of peritoneal macrophages

The difference in macrophage replenishment prompted us to assess the wider effects of sex on the imprinting of peritoneal macrophage identity and function. We therefore first performed population-level RNASeq on peritoneal F4/80^hi^CD102^+^ macrophages FACS-purified from unmanipulated 10-12 week old male and female mice (**Supplementary Figure 4**). To limit potential confounding effects of estrous cycle, the stage of each female mouse was confirmed by vaginal cytology and samples pooled to include cells from all stages of the cycle. Furthermore, to limit the effects of circadian influence, mice in each biological replicate were euthanised at the same time each day. Unbiased clustering was then used to group the populations based on sex, with sex explaining 81% of the variance within the datasets (Figure 4a) and differential gene expression analysis revealed that 486 mRNA transcripts were differentially expressed (>1.5fold) between female and male peritoneal CD102^+^ macrophages (Figure 4b **& Supplementary Table 1**). Analysis of the 148 mRNA transcripts more highly expressed in female peritoneal macrophages revealed that a large proportion was associated with immune function, including the C-type lectin receptors *Clec4g*, *Cd209a* and *Cd209b*, the complement components *C4b*, *C1qa*, and *C3* the immunoregulatory cytokine *Tgfb2*, the B cell chemoattractant *Cxcl13*, and as expected the phagocytic receptor *Timd4* (Figure 4c **& Supplementary Table 1**). Consistent with this, ‘immune response’ and ‘immune system processes’ were among the top pathways identified by gene-set enrichment analysis in genes up-regulated in female cells (**Supplementary Table 2**). Transcripts for the apolipoproteins *Apoe, Saa2, Saa3* and *Apoc1* were also expressed more highly in female cells. Notably, in contrast to previous work that assessed basal gene expression by total peritoneal cells across the sexes ^38^, we did not detect any dimorphism in expression of toll-like receptors (TLRs), the TLR adaptor molecule MyD88 or CD14 (**Supplementary Figure 5**). Moreover, the dimorphic cassette of genes we identified is distinct from that recently shown to be sexually dimorphic in microglia (**Supplementary Figure 5**).

**Figure 4:**
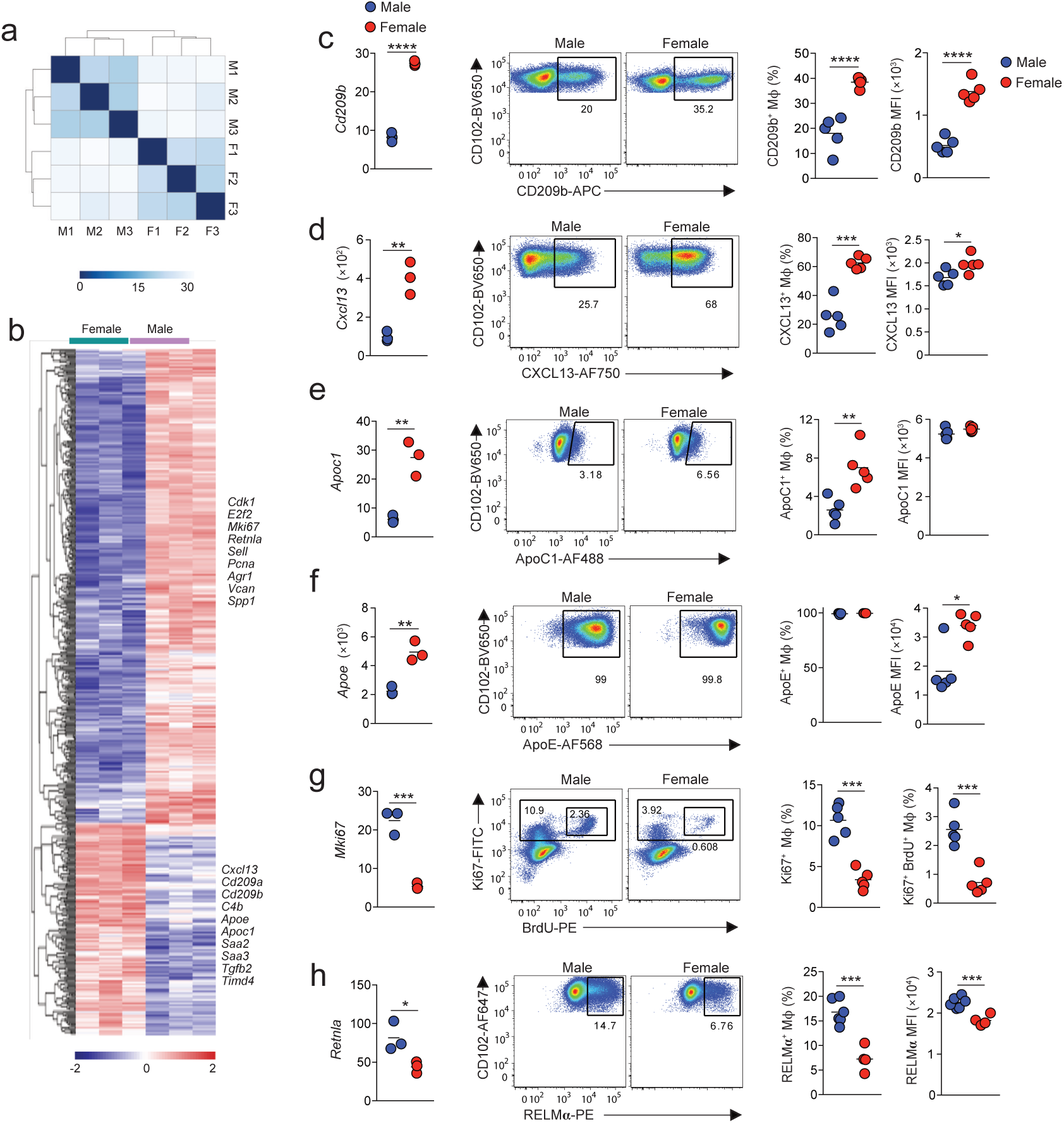
Sex determines the transcriptional signature of peritoneal macrophages. (**a**) Heatmap showing distance between samples of male (M) and female (F) CD102^+^F4/80^hi^ macrophages FACS-purified from the peritoneal cavity of 10-12 week old mice. (**b**) Gene expression profile of the 148 differentially expressed (>1.5 fold) genes between male and female peritoneal macrophages with selected genes highlighted. (**c**) Expression of *Cd209b* from RNAseq (FPKM; *left panel*), representative expression of CD209b protein (*middle panels*) and frequency of CD209b^+^ cells amongst CD102^+^F4/80^hi^ peritoneal macrophages obtained from 10-12-week-old male or female C57BL/6 mice (*right panel*) and the mean fluorescence intensity (MFI) of CD209b expression by these cells (*far right panel*). Symbols represent individual animals and horizontal bars represent the mean. RNAseq data represent 3 mice per group and protein analysis represents 5 mice per group from one of five independent experiments. ****P<0.0001. Student’s *t* test. (**d**) Expression of *Cxcl13* from RNAseq (FPKM; *left panel*), representative expression of CXCL13 mRNA (*middle panels*) and frequency of CXCL13^+^ cells amongst CD102^+^F4/80^hi^ peritoneal macrophages obtained from 10-12-week-old male or female C57BL/6 mice (*right panel*) and the mean fluorescence intensity (MFI) of CXCL13 mRNA expression by these cells (*far right panel*). Symbols represent individual animals and horizontal bars represent the mean. RNAseq data represent 3 mice per group and flow cytometric analysis represents 5 mice per group from one of three independent experiments. *P<0.05, ***P<0.001. Student’s *t* test. (**e**) Expression of *Apoc1* from RNAseq (FPKM; *left panel*), representative expression of ApoC1 mRNA (*middle panels*) and frequency of ApoC1^+^ cells amongst CD102^+^F4/80^hi^ peritoneal macrophages obtained from 10-12-week-old male or female C57BL/6 mice (*right panel*) and the mean fluorescence intensity (MFI) of ApoC1 mRNA expression by these cells (*far right panel*). Symbols represent individual animals and horizontal bars represent the mean. RNAseq data represent 3 mice per group and flow cytometric analysis represents 5 mice per group from one of three independent experiments. *P<0.05, ***P<0.001. Student’s *t* test. (**f**) Expression of *Apoe* from RNAseq (FPKM; *left panel*), representative expression of ApoE mRNA (*middle panels*) and frequency of ApoE^+^ cells amongst CD102^+^F4/80^hi^ peritoneal macrophages obtained from 10-12-week-old male or female C57BL/6 mice (*right panel*) and the mean fluorescence intensity (MFI) of ApoE mRNA expression by these cells (*far right panel*). Symbols represent individual animals and horizontal bars represent the mean. RNAseq data represent 3 mice per group and flow cytometric analysis represents 5 mice per group from one of three independent experiments. *P<0.05, ***P<0.001. Student’s *t* test. (**g**) Expression of *Mki67* from RNAseq (FPKM; *left panel*), representative expression of Ki67 protein and BrdU incorporation (*middle panels*) and the frequency of BrdU^+^Ki67^+^ cells amongst CD102^+^F4/80^hi^ peritoneal macrophages obtained from 10-12-week-old male or female C57BL/6 mice. Symbols represent individual animals and horizontal bars represent the mean. Data represent 5 mice per group from one of two experiments. ***P<0.001. Student’s *t* test. (**h**) Expression of *Retnla* from RNAseq (FPKM; *left panel*), representative expression of RELMα protein (*middle panels*) and the frequency of RELMα^+^ cells amongst CD102^+^F4/80^hi^ peritoneal macrophages obtained from 10-12-week-old male or female *Rag1*^−/−^ C57BL/6 mice. Symbols represent individual animals and horizontal bars represent the mean. *P<0.05, ***P<0.001. Student’s *t* test.

We used flow cytometry to confirm higher expression of *Cd209b*, *Cxcl13*, and *Apoc1* by female macrophages, as these were the most differentially expressed non-X-linked genes with mapped read counts greater than 10. This analysis revealed unexpected heterogeneity within resident peritoneal macrophages. For instance, only a proportion of male and female CD102^+^ macrophages expressed CD209b, although the frequency of these was greater in females than in males (35% and 20% respectively). Moreover, CD209b was expressed at a higher level on a per cell basis by female CD102^+^ macrophages compared with their male counterparts (Figure 4c), a finding consistent across different strains, including *Rag1*^−/−^ mice, and mice from different housing environments (**Supplementary Figure 6**). Due to the unavailability of commercial antibodies for CXCL13 and ApoC1, we used PrimeFlow technology to measure CXCL13 and ApoC1 mRNA at a single cell level using flow cytometry. Again, this revealed that a greater proportion of female CD102^+^ macrophages expressed mRNA for CXCL13 and ApoC1 than their male counterparts, and CXCL13 mRNA was also higher on a per cell basis in female cells (Figure 4d, e). In contrast, PrimeFlow measurement of mRNA for ApoE, the most highly expressed of all differentially expressed genes by female cells by RNAseq, revealed that all peritoneal macrophages expressed ApoE irrespective of sex, but that expression was higher in female cells on a per cell basis. Hence, the transcriptional differences seen at population level appear to result from differential gene expression at a single cell level but also from different frequencies of gene-expressing cells amongst the CD102^+^ population.

The majority of genes more highly expressed by male peritoneal CD102^+^ macrophages were associated with cell cycle, including *Cdk1, E2f2,* and *Mki67* (Figure 4b **& Supplementary Table 1**). Pathway analysis also revealed that at least 162 of the 338 genes differentially up-regulated in male CD102^+^ macrophages were associated with proliferation, and cell cycle-related processes predominated among the significantly enriched pathways (**Supplementary Table 2)**. Short-term BrdU pulse-chase experiments confirmed that male CD102^+^ macrophages have elevated levels of *in situ* proliferation compared with their female counterparts (Figure 4g). These analyses also identified that *Retlna,* which encodes the immunomodulatory cytokine RELMα and is expressed specifically by those resident peritoneal macrophages that are most recently-derived from monocytes ^21^, was differentially expressed between sexes, with higher expression by male cells at both the mRNA and protein level (Figure 4h).

Of note, a number of genes previously reported to distinguish long-lived, embryonically-derived macrophages from those of recent BM origin in the lung and liver were more highly expressed in females. These included receptors involved in phagocytosis and immunity (i.e. *Timd4*, *Colec12*, and *Cd209* family members), *Apoc1*, as well as the bone morphogenic receptor *Bmpr1a* (**Supplementary Table 1**). Lowering the stringency of selection of differentially-expressed genes identified additional genes within the female-specific cluster that have been associated with embryonically-derived or long-lived macrophages, including *Marco* and *Cd163* that also encode phagocytic receptors (data not shown). Furthermore, to discern systemic from local effects of sex, we compared gene expression by CD102^+^ macrophages from female peritoneal cavity to pleural CD102^+^ macrophages from both sexes. This analysis identified a module of 18 genes that was uniquely upregulated by female peritoneal macrophages, and that included *Apoc1*, *Cd209b*, and *Colec12*, as well as *Saa3*, *C4b*, and *Tgfb2* (**Supplementary Table 3**). Conversely, the 86 genes uniquely downregulated by female peritoneal macrophages compared with the other CD102^+^ populations were highly enriched for cell cycle related genes and pathways (**Supplementary Table 3&4**). Thus, the more limited proliferative activity of female peritoneal macrophages and their expression of numerous immune-related genes appear either related to their slower replenishment from the bone-marrow or regulated directly by the unique signals present within the female peritoneal microenvironment.

### scRNAseq analysis reveals dimorphic macrophage heterogeneity

We next applied single cell RNA sequencing (scRNAseq) to determine whether the transcriptional differences seen in our population-level data were the result of gene differences at a single cell level or if dimorphism was a reflection of differential subset composition between the sexes. A broad approach was used to capture all CD11b^+^ cells depleted of granulocytes and B1 B cells to allow both CD102^−^ F4/80^lo^MHCII^+^ macrophages and resident CD102^+^ cells to be examined. These cells were FACS-purified from age-matched 12-week-old male and female mice and droplet-based scRNASeq performed using the 10X Genomics platform. 10,000 sorted cells of each sex were sequenced and following quality control, analysis was performed on 4341 and 2564 cells from female and male respectively.

Uniform Manifold Approximation and Projection (UMAP) dimensionality reduction analysis revealed 6 clusters that were present in both male and female cells (Figure 5a, b). Given that the starting population of CD11b^+^ cells is known to be phenotypically heterogeneous, containing resident CD102^+^F4/80^hi^ macrophages, CD102^−^F4/80^lo^ MHCII^+^ macrophages and CD11c^+^MHCII^+^ cDC2 ^20–23^, we first used a panel of known markers to validate subset identity (Figure 5b). 3 clusters of resident macrophages (3-5) could be identified on the basis of their high expression of *Adgre1* (F4/80) and *Icam2* (CD102). As expected, these were clearly distinct from short-lived CD102^−^F4/80^lo^MHCII^+^ macrophages and cDC2, which were found in clusters 1 and 2 respectively, and expressed *Ccr2* (Figure 5c, d) ^21^. However, CD102^−^F4/80^lo^MHCII^+^ macrophages and cDC2 could be distinguished from one another on the basis of expression of the DC markers *Cd209a* and *Napsa* ^27, 43^, and of *Retnla* and *Fclrs*, which we and others have shown to be signature markers of cavity CD102^−^F4/80^lo^MHCII^+^ macrophages ^21, 22, 44^. Cluster 6 was defined by genes associated with cell cycle, such as *Mki67* and *Birc5*, suggesting this cluster represents proliferating cells. In both sexes, the majority of cells was in cluster 5 (Figure 5e), which was characterised by markers of resident F4/80^hi^CD102^+^ macrophages including *Icam2*, *Prg4* and *Tgfb2* (Figure 5c, d **& Supplementary Table 5)** that form part of the core peritoneal macrophage-specific transcriptional signature ^45^; cluster 5 cells also expressed markers of long-lived macrophages, including *Timd4* and *Apoc1*, confirming the findings above. Although the cells in cluster 3 expressed *Icam2*, they also expressed a number of genes that were highly expressed by the CD102^−^ F4/80^lo^MHCII^+^ macrophages in cluster 2, such as *Retnla, H2.Aa* and *Ccr2*, suggesting a common origin of these clusters, or a close relationship between them. This analysis also identified genes uniquely expressed by cluster 3, including *Folr2*, which encodes the beta subunit of the folate receptor (FRβ). Although cluster 4 showed a distinct pattern of gene expression, such as high expression of *Apoe*, it also shared features with cluster 3 and cluster 5, suggesting it may contain differentiation intermediates. Consistent with our earlier analysis, we found that the *Timd4*-expressing cluster 5 was more abundant amongst female cells, whereas more male cells were found within clusters 1, 2, 3 and 6 (Figure 5c).

**Figure 5:**
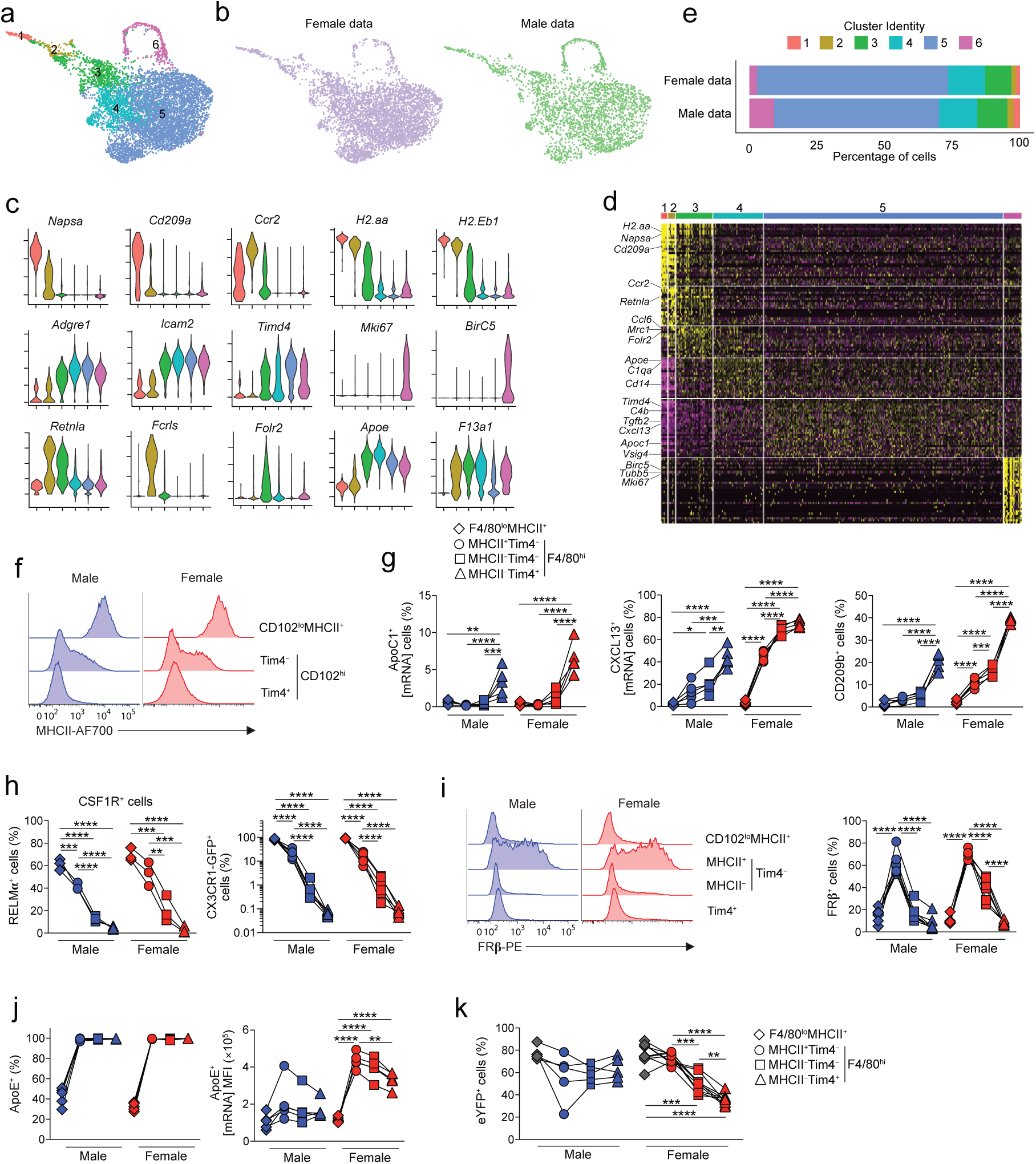
scRNAseq analysis reveals dimorphic macrophage heterogeneity. (**a**) UMAP dimensionality reduction analysis of 4341 and 2564 number of cells from the peritoneal cavity of 19 week-old male or female mice identifying 6 clusters. (**b**) UMAP profile of female and male peritoneal cells. (**c**) Feature plots displaying expression of individual genes by merged female/male cells. (**d**) Heatmap displaying the 10 most differentially expressed genes by each cluster from **a** (select genes highlighted). (**e**) Relative frequency of each cluster in the female and male dataset. (**f**) Representative expression of MHCII by CD102^lo^MHCII^+^ and Tim4/MHCII-defined CD102^+^ peritoneal macrophages from 10-12 week old male or female C57BL/6 mice. (**g**) Expression of Apoc1 (mRNA), CXCL13 (mRNA) and CD209b protein by CD102^lo^MHCII^+^ and Tim4/MHCII-defined CD102^+^ peritoneal macrophages from 10-12 week old male or female C57BL/6 mice. Data represent 5 mice per group from one of three independent experiments. *P<0.05, **P<0.01, ***P<0.001, ****P<0.0001. One-way ANOVA with Tukey’s multiple comparisons test. (**h**) Expression of RELMα and CX3CR1-GFP by CD102^lo^MHCII^+^ and Tim4/MHCII-defined CD102^+^ peritoneal macrophages from 10 week old (RELMa) or 15 week old (CX3CR1-GFP) male or female C57BL/6 mice. Data represent 3 (RELMα), 5 (CX3CR1-GFP, female) or 7 (CX3CR1-GFP, male) mice per group from one of at least 5 independent experiments (RELMα) or from one experiment (CX3CR1-GFp). *P<0.05, **P<0.01, ***P<0.001, ****P<0.0001. One-way ANOVA with Tukey’s multiple comparisons test. (**i**) Histograms show representative expression of FRβ by CD102^lo^MHCII^+^ and Tim4-defined CD102^+^ peritoneal macrophages from 10-12 week old male or female C57BL/6 mice. Scatter plot show frequency of FRβ^+^ cells amongst CD102^lo^MHCII^+^ and Tim4/MHCII-defined CD102^+^ peritoneal macrophages. Data represent 6 (female) or 7 (male) mice per group pooled from two independent experiments. *P<0.05, **P<0.01, ***P<0.001, ****P<0.0001. One-way ANOVA with Tukey’s multiple comparisons test. (**j**) Frequency of ApoE^+^ (mRNA) cells amongst CD102^lo^MHCII^+^ and Tim4/MHCII-defined CD102^+^ peritoneal macrophages (*left*) and mean fluorescence intensity of ApoE by these subsets (*right*) from 10-12 week old male or female C57BL/6 mice. Data represent 5 mice per group pooled from one of 3 independent experiments. *P<0.05, **P<0.01, ***P<0.001, ****P<0.0001. One-way ANOVA with Tukey’s multiple comparisons test. (**k**) Frequency of eYFP^+^ cells amongst F4/80, MHCII and Tim4-defined macrophages obtained from 16-week-old male and female CD11c^Cre^.*Rosa26*^LSL-eYFP^ mice. Symbols represent individual animals and horizontal bars represent the mean. Data represent 5 (male) or 9 (female) mice per group pooled from two independent experiments.

We next used flow cytometry to determine if we could validate the additional heterogeneity uncovered by our scRNAseq analysis. Given that MHCII-associated genes appeared to define heterogeneity amongst Tim4^−^CD102^+^ macrophages (i.e. clusters 3 & 4), we assessed expression of MHCII by Tim4-defined subsets of CD102^+^ macrophages. This confirmed that a proportion of Tim4^−^ CD102^+^ macrophages expressed MHCII, albeit at lower levels than CD102^−^F4/80^lo^MHCII^+^ macrophages, whereas Tim4^+^ macrophages had negligible MHCII expression (Figure 5f). Thus, we used a combination of Tim4 and MHCII to identify macrophage subsets and assessed expression of other subset defining markers from the scRNAseq analysis. Consistent with the analysis above, we found that ApoC1 expression was essentially exclusive to Tim4^+^MHCII^−^ macrophages (Figure 5g), whereas both MHCII-defined Tim4^−^ macrophages lacked ApoC1 expression. Expression of CXCL13 was also highest amongst Tim4^+^MHCII^−^ cells, although interestingly, the proportion of CXCL13^+^ cells increased progressively from Tim4^−^MHCII^+^ to Tim4^−^MHCII^−^ to Tim4^+^MHCII^−^ CD102^+^ macrophages (Figure 5g). Although not identified as a cluster defining gene in our scRNAseq analysis due to low coverage, we found that CD209b displayed the same pattern of expression as CXCL13 (Figure 5g). Consistent with the idea that they may derive from CD102^−^F4/80^lo^MHCII^+^ macrophages, the expression of RELMα and CX3CR1 was highest on Tim4^−^MHCII^+^ macrophages and was essentially absent from Tim4^+^MHCII^−^ CD102^+^ macrophages. The majority of Tim4^−^MHCII^+^ macrophages expressed high levels of FRβ, whereas all other populations had little or no expression, consistent with our scRNAseq analysis. While *Apoe* was proposed to define cluster 4 in our scRNAseq analysis, consistent with our analysis above, we found it was expressed by all CD102^+^ macrophages, although, in females, most highly expressed by Tim4^−^MHCII^+^ and the level decreased progressively to Tim4^+^MHCII^−^ CD102^+^ macrophages. Finally, we returned to using CD11c^Cre^.*Rosa26*^LSL-eYFP^ mice to assess if MHCII/Tim4-defined subsets showed differential levels of replenishment. Notably, we found that MHCII-expressing Tim4^−^CD102^+^ macrophages showed equivalent labelling to CD102^−^F4/80^lo^MHCII^+^ macrophages in female CD11c^Cre^.*Rosa26*^LSL-eYFP^ mice, indicative of more recent derivation from CD102^−^ F4/80^lo^MHCII^+^ cells. Consistent with their intermediate transcriptional profile, Tim4^−^CD102^+^ macrophages that had lost MHCII expression showed intermediate labelling when compared with their MHCII^+^Tim4^−^ and Tim4^+^ counterparts. No difference in eYFP labelling between Tim4-defined subsets was noted in male mice, consistent with more rapid replenishment of all subsets of macrophages in this environment.

Collectively these data show that excluding proliferating cells, resident peritoneal macrophages comprise three main clusters, with Tim4^−^ macrophages displaying an intermediate phenotype compared with F4/80^lo^MHCII^+^ macrophages and Tim4^+^ macrophages.

### Differential replenishment and environmental signals drive the dimorphic features of peritoneal macrophages

To dissect the dimorphic features of CD102^+^ macrophages that could be related to longevity from those more directly controlled by dimorphic environmental signals, we next assessed expression of these in *Ccr2^−/−^* mice in whom macrophage replenishment is markedly reduced due to severe monocytopenia ^46, 47^. Strikingly, the frequency of Tim4^−^ macrophages, as well as those expressing RELMα, FRβ or MHCII were markedly reduced in *Ccr2^−/−^* mice compared with *Ccr2^+/+^* mice irrespective of sex, confirming these cells to be recently derived from monocytes (Figure 6a **& Supplementary Figure 7**). In males, CCR2 deficiency also led to reduced expression of ApoE and emergence of an ApoE^−^ subset of CD102^+^ macrophages (Figure 6b). In contrast, a higher proportion of CD102^+^ macrophages in *Ccr2^−/−-^* mice expressed CD209b and ApoC1, markers that are characteristic of the Tim4^+^MHCII^−^ subset, suggesting these markers may be expressed selectively by long-lived macrophages (Figure 6b). Consistent with this, Tim4^+^ macrophages expressing CD209b displayed the lowest level of replacement by donor cells in BM chimeras when compared with all other CD209b/Tim4-defined macrophages, even in male mice where overall replenishment from the bone marrow is markedly higher (Figure 6c). The low levels of replacement of peritoneal CD209b^+^Tim4^+^ macrophages does not reflect derivation from yolk sac progenitors, as, unlike microglia in the brain, these cells are not labelled in male or female *Cdh5*^Cre-ERT2^.*Rosa*^LSL*-*tdT^ mice, which allow tracing of cells arising from yolk sac haematopoiesis ^48^. Similar results were obtained with CD209b^+^Tim4^+^ macrophages in the pleural cavity (Figure 6d). Hence, despite being long-lived, CD209b^+^Tim4^+^ macrophages derive from conventional haematopoiesis in both sexes. Importantly, temporal analysis revealed that while little difference in abundance of CD209b-expressing CD102^+^ macrophages was seen in pre-pubescent (4-5-week-old) male and female mice, these cells accumulated progressively in the cavity of female mice following sexual maturation. This did not occur in male mice, consistent with their higher rate of replenishment from the bone marrow and indicating that acquisition of CD209b expression appears to be associated with time-of-residency in the female cavity (Figure 6e).

**Figure 6:**
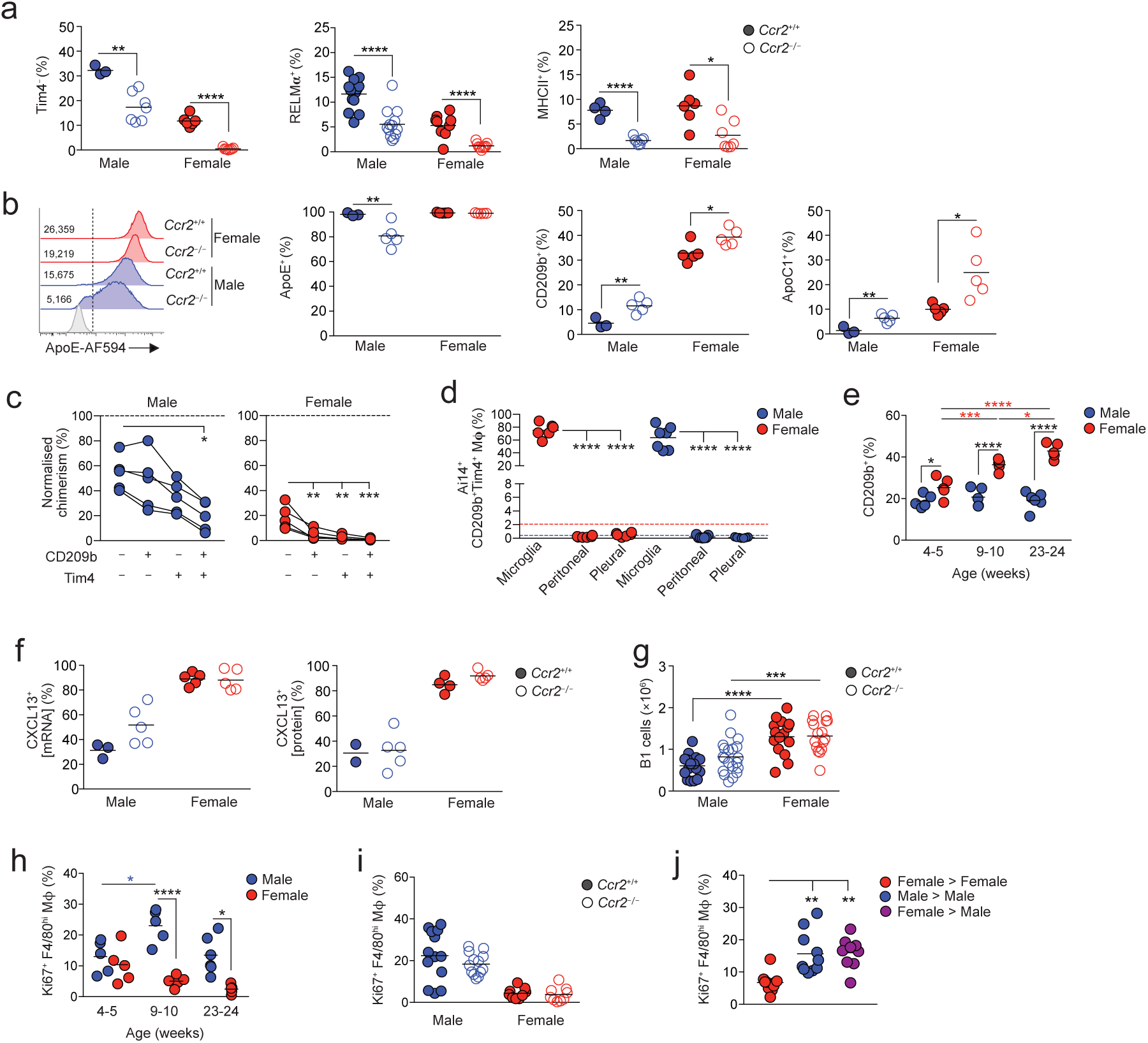
Differential replenishment and environmental signals drive the dimorphic features of peritoneal macrophages. (**a**) Frequency of Tim4^−^, RELMα^+^ and MHCII^+^ cells amongst CD102^+^ macrophages from the peritoneal cavity of unmanipulated age-matched *Ccr2*^+/+^ or *Ccr2*^−/−^ mice. Symbols represent individual animals and horizontal bars represent the mean. Data are pooled from two independent experiments. Tim4 data represents 3 (*Ccr2*^+/+^ males), 6 (*Ccr2*^+/+^ females) or 7 (*Ccr2*^−/−^) 22-28 week old mice per group. RELMα data represent with 13 male and 9 female 14-18 week old mice per group. MHCII data represent 4 (*Ccr2*^+/+^ males), 6 (*Ccr2*^+/+^ females) or 7 (*Ccr2*^−/−^) 22-28 week old mice per group. *P<0.05, **P<0.01, ****P<0.0001. Student’s *t* test with Holm-Sidak correction. (**b**) Representative expression of ApoE by CD102^+^ macrophages (*histograms*) and frequency of ApoE^−^, CD209b^+^ and ApoC1^+^ cells from the peritoneal cavity of unmanipulated age-matched *Ccr2*^+/+^ or *Ccr2*^−/−^ mice. Symbols represent individual animals and horizontal bars represent the mean. Data are pooled from two independent experiments and represents 3 (*Ccr2*^+/+^ males), 6 (*Ccr2*^+/+^ females) or 7 (*Ccr2*^−/−^) 22-28 week old mice per group. *P<0.05, **P<0.01. Student’s *t* test with Holm-Sidak correction. (**c**) Normalized non-host chimerism of CD209/Tim4-defined subsets of CD102^+^ macrophages from the peritoneal cavity of sex matched tissue-protected BM chimeric mice 8 weeks post-reconstitution. Data are normalised to the non-host chimerism of Ly6C^hi^ monocytes. Data represent 5 mice per group from one experiment. *P<0.05, ***P<0.001. Paired Student’s *t* test. (**d**) Proportion of tdTomato^+^ (Ai14) cells amongst microglia, peritoneal and pleural macrophages from 15-week-old *Cdh5*^Cre-ERT2^.*Rosa26*^LSL-Ai14^.*Cx3cr1*^+/gfp^ mice administered 4-hydroxytamoxifen at E7.5. Data represent 6 (female) or 7 (male) mice per group from one experiment. ****P<0.0001. One-way ANOVA followed by Tukey’s multiple comparisons test. (**e**) Frequency of cells expressing CXCL13 mRNA (*left*) or CXCL13 protein (*right*) amongst CD102^+^ macrophages obtained from the peritoneal cavity of unmanipulated 22-28 week old *Ccr2*^+/+^ or *Ccr2*^−/−^ mice. Symbols represent individual animals and horizontal bars represent the mean. CXCL13 mRNA data represents 3 (*Ccr2*^+/+^ males), 5 (*Ccr2*^+/+^ females) or 5 (*Ccr2*^−/−^) mice per group. CXCL13 protein data represents 2 (*Ccr2*^+/+^ males), 4 (*Ccr2*^+/+^ females) or 5 (*Ccr2*^−/−^) mice per group. (**f**) The absolute number of B1 cells obtained from the peritoneal cavity of unmanipulated age matched 14-28 week old *Ccr2*^+/+^ or *Ccr2*^−/−^ mice. Data represent 15 (*Ccr2*^+/+^ females), 16 (*Ccr2*^−/−^ females), 17 (*Ccr2*^+/+^ males) or 20 (*Ccr2*^−/−^ females) mice per group pooled from four independent experiments. (**g**) Frequency of Ki67^+^ cells amongst peritoneal F4/80^hi^ macrophages obtained from the peritoneal cavity of unmanipulated 14-18 week old *Ccr2*^+/+^ or *Ccr2*^−/−^ mice. Data represents 15 (*Ccr2*^+/+^ females), 16 (*Ccr2*^−/−^ females), 17 (*Ccr2*^+/+^ males) or 20 (*Ccr2*^−/−^ females) mice per group pooled from 2 experiments. (**h**) Frequency of Ki67^+^ cells amongst peritoneal F4/80^hi^ macrophages obtained from sex matched or mismatched tissue protected BM chimeric mice 8-12 weeks post-reconstitution. Data represent 9 (female > male) or 10 (sex matched groups) mice per group pooled from one of two independent experiments. **P<0.01. One-way ANOVA followed by Tukey’s multiple comparisons test.

Not all dimorphic features of peritoneal CD102^+^ macrophages were influenced by their rate of replenishment. For instance, the intrinsically higher expression of CXCL13 by female macrophages was not altered by CCR2 deficiency (Figure 6f). In parallel, although we confirmed previous findings of a clear dimorphism in the numbers of B1 cells between adult male and female mice ^15^ and this developed gradually following sexual maturation (**Supplementary Figure 8**), this phenomenon remained in *Ccr2^−/−^* mice (Figure 6g). Similarly, while the higher levels of proliferation by male CD102^+^ macrophages developed following sexual maturation (Figure 6h), this was unaffected by CCR2 deficiency (Figure 6i). This evidence that certain dimorphic features are driven by environmental factors, independent of cell replenishment was supported further by the fact that macrophages derived from female BM in the cavity of chimeric male showed levels of proliferation that were identical to those of male BM derived macrophages in the male cavity and were higher than those of female BM derived macrophages in female cavity (Figure 6i). Thus, the differential proliferation of female and male macrophages is not due to cell-intrinsic differences in their proliferative activity.

Taken together these data demonstrate that both local imprinting and differential turnover contribute to the sexual dimorphisms seen in peritoneal macrophages.

### Differential CD209b expression confers an advantage on female macrophages in the setting of pneumococcal peritonitis

We postulated that differential expression of key pattern recognition receptors such as CD209b might endow female macrophages with an enhanced ability to deal with bacterial infection. To test this idea, we examined the acute peritonitis caused by the gram-positive bacterium *Streptococcus pneumoniae* (Figure 7a), a localised model of infection in which resident macrophages, and in particular CD209b, are indispensable for protective immunity ^49, 50^ whereas recruitment of neutrophils is not required ^49^and hence avoids any confounding effects of systemic sex-dependent effects on innate immune responses that have been reported previously ^51^. As CD209b is expressed exclusively by CD102^+^ macrophages in the peritoneal cavity (**Supplementary Figure 6**), this model allowed us to directly assess the importance of differential CD209b expression by CD102^+^ macrophages in bacterial elimination. Strikingly, females showed enhanced capability to control *S. pneumoniae* infection, with significantly lower levels of bacteria in peritoneal fluid of female mice compared with their male counterparts 20hrs after inoculation (Figure 7b). Fewer neutrophils and Ly6C^hi^ monocytes were present in the female cavity compared with male mice (Figure 7c), consistent with a model in which resident macrophages control infection ^49^. In contrast, while the well-documented macrophage ‘disappearance reaction’ ^52^ occurred in both male and female mice after infection (Figure 7d), significantly higher numbers of CD209b-expressing macrophages persisted in the female cavity (Figure 7d). Administration of an anti-CD209b blocking antibody (22D1) ^53^ led to increased levels of bacteraemia in female mice, although this did not attain statistical significance due to variance in bacterial counts in some mice in whom the macrophage ‘disappearance reaction’ was more pronounced (Figure 7e).

**Figure 7:**
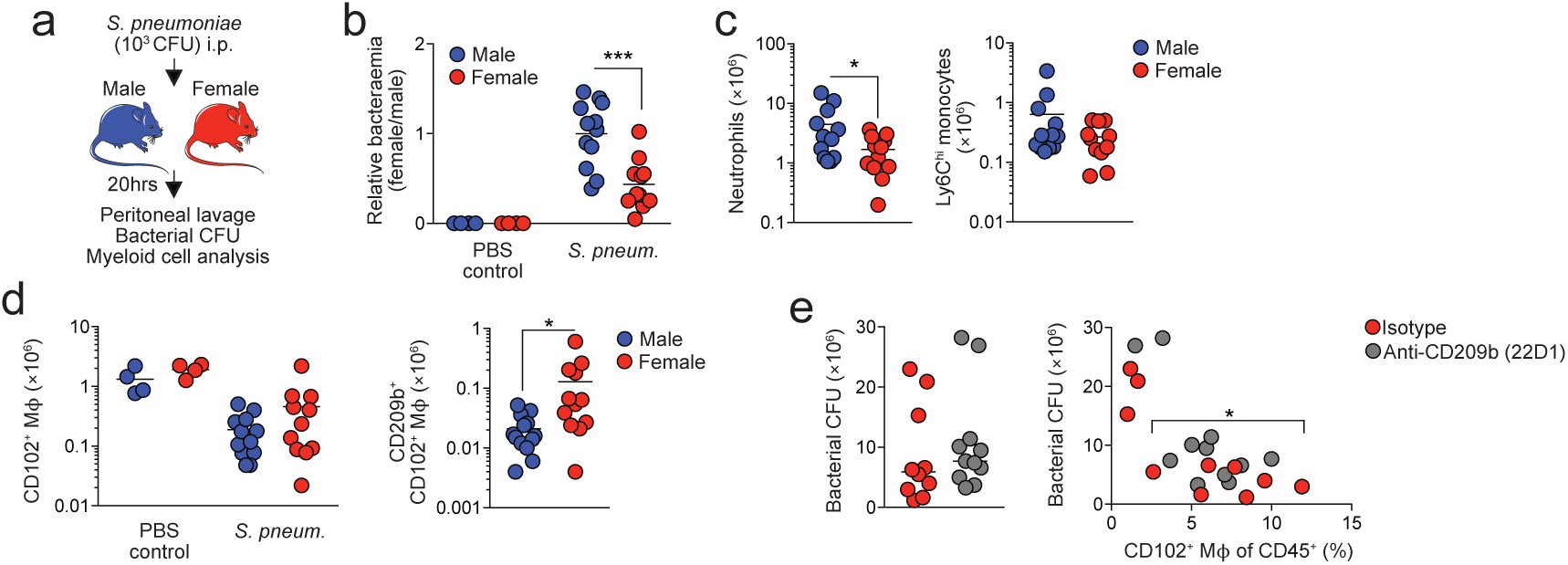
Differential CD209b expression confers an advantage on female macrophages in the setting of pneumococcal peritonitis. (**a**) Experimental scheme for induction of peritonitis. Male and female mice (9-10 weeks old) were inoculated with 10^3^ CFU type 2 *Streptococcus pneumoniae* (D39) and bacterial counts and assessment of the peritoneal myeloid compartment assessed after 20hrs. (**b**) Relative bacteraemia in the peritoneal cavity of male and female mice infected 20hrs earlier (female CFU/male CFU). Symbols represent individual animals and horizontal bars represent the mean. Data represent 4 (PBS), 11 (female *S. pneumoniae*) or 12 (male *S. pneumoniae*) mice per group pooled from three independent experiments. ***P<0.001. Student’s *t* test. (**c**) Absolute numbers of Ly6G^+^ neutrophils and Ly6C^hi^ monocytes in the peritoneal cavity 20hrs after inoculation with 10^3^ CFU type 2 *Streptococcus pneumoniae* (D39) or in mice that received PBS. Symbols represent individual animals and horizontal bars represent the mean. Data represent 4 (PBS), 11 (female *S. pneumoniae*) or 12 (male *S. pneumoniae*) mice per group pooled from three independent experiments. *P<0.05. Student’s *t* test. (**d**) Absolute numbers of CD102^+^ macrophages and CD209b^+^CD102^+^ macrophages in the peritoneal cavity 20hrs after inoculation with 10^3^ CFU type 2 *Streptococcus pneumoniae* (D39) or in mice that received PBS. Symbols represent individual animals and horizontal bars represent the mean. Data represent 4 (PBS), 11 (female *S. pneumoniae*) or 12 (male *S. pneumoniae*) mice per group pooled from three independent experiments. *P<0.05. Student’s *t* test. (**e**) Numbers of bacteria in the peritoneal cavity of male and female mice infected 20hrs earlier and pre-treated with anti-CD209b (22D1) or a matched isotype control (Ham IgG1) 30mins prior to inoculation (*left*). Right, bacteraemia versus the frequency of CD102^+^ macrophages in the peritoneal cavity of mice above. Symbols represent individual animals and horizontal bars represent the mean. Data represent 10 (isotype control) or 11 (anti-CD209b) mice per group pooled from two independent experiments.

Thus, dimorphic expression of key immune receptors and molecules leads to differential ability to handle local bacterial infection.

## Discussion

Understanding the extrinsic and intrinsic factors that govern tissue macrophage differentiation is a key goal in the field of macrophage biology. Here we reveal a striking effect of sex on the phenotypic and transcriptional identity of resident peritoneal macrophages and demonstrate that this contributes to the sex-dependent resistance of mice to bacterial peritonitis. Moreover, we show that this arises through a combination of dimorphic microenvironmental signals and sex-dependent differences in the rate of macrophage renewal from the bone marrow.

Using classical defining markers such as F4/80, CD11b and CD102, we found peritoneal macrophages from male and female mice to be phenotypically identical. Furthermore, while some studies have reported that the number of peritoneal macrophages is greater in females ^38, 51^, we did not routinely detect significant differences in the number of peritoneal macrophages between the sexes. However, mRNA sequencing revealed marked dimorphism in the transcriptional fingerprint of resident peritoneal macrophages under homeostatic conditions. Importantly, this showed that female CD102^+^ macrophages express higher levels of genes associated with lipid uptake and transport as well as immune defence/response, including those encoding complement components, the chemoattractant CXCL13, and numerous receptors involved in recognition and uptake of pathogens and apoptotic cells. In contrast, the signature of male peritoneal macrophages was dominated by cell cycle associated genes consistent with their elevated levels of proliferation, a dimorphism we have reported previously ^21^. Although others have reported dimorphic expression of TLRs and CD14 by peritoneal macrophages, no consistent pattern was observed in these studies ^37, 38^ and we found no significant difference in mRNA transcripts of the adaptor protein MyD88 or any TLRs, consistent with more recent analysis of surface protein expression ^51^. We have also found no sex differences in CD14 expression at the gene or protein level. These discrepancies may relate to differences in the nature of the cells being analysed, with one study using peritoneal macrophages elicited by incomplete Freund’s adjuvant ^37^ and the other assessing gene expression by total peritoneal cells ^38^. In contrast, our analyses involved minimal handling and used rigorously characterized resident macrophages.

Our further analyses revealed that many of the sexually dimorphic features of macrophages arose following sexual maturation, including the higher expression of CD209b and Tim4 by female macrophages. Furthermore, the enhanced accumulation of peritoneal B1 cells found in females was also age-dependent, suggesting that the higher levels of CXCL13 production by female macrophages may also be driven by sexual maturation. Dimorphic differences in the turnover of CD102^+^ resident peritoneal macrophages also appeared to largely arise following sexual maturation. Notably, the kinetics of labelling in CD11c^cre^.*Rosa26*^LSL-eYFP^ mice and a significant reversal in the autonomy of female peritoneal macrophages in BM chimeras following ovariectomy suggest this aspect of dimorphism in normal mice reflects a switch from replenishment to self-maintenance in females. These results are consistent with recent monocyte fate mapping using *Ms4a3*^Cre^.*Rosa26*^LSL-tdTomato^ mice showing that while monocytes contribute to the maintenance of peritoneal macrophages during the perinatal and adolescent period, this process wanes during adulthood in female mice (^54^ & F. Ginhoux, personal communication). Hence, sexual maturation leads to dimorphic changes in replenishment, proliferation and gene expression by peritoneal macrophages, at least some of which are driven by the female peritoneal environment.

Despite a significant degree of transcriptional difference at the population level, single cell mRNA sequencing showed that male and female resident macrophages encompassed very similar transcriptionally-defined clusters of cells. However, the relative abundance of these clusters differed between sexes. In this regard, CD102^+^ peritoneal macrophages could be divided into three predominant transcriptionally-distinct clusters. Of these, the cells in cluster 3 expressed CD102 together with MHCII, RELMα and CX3CR1, all of which are key markers of F4/80^lo^MHCII^+^ peritoneal macrophages, suggesting that cluster 3 may be recently derived from the F4/80^lo^MHCII^+^ macrophage population that is derived from blood monocytes in adult mice ^21–23^. As cells in cluster 4 shared features with both the MHCII-defined cluster 3 and the dominant cluster 5 population of Tim4^+^ macrophages, these may represent a further intermediate differentiation state. Consistent with this idea, in female mice Tim4^−^MHCII^−^ CD102^+^ macrophages, which likely represent those in cluster 4, displayed an intermediate degree of replenishment from the bone marrow in our fate mapping studies between that of the rapidly replenished Tim4^−^MHCII^+^ CD102^+^ macrophages and slowly replenished Tim4^+^MHCII^−^ cells. Furthermore, all Tim4^−^ cells were largely ablated in female *Ccr2*^−/−^ mice. A linear-developmental relationship that culminates at cluster 5 would be consistent with the greater abundance of cells in this cluster in females, given the slower entry of bone-marrow-derived cells into the female CD102^+^ macrophage pool. However, it seems unlikely that such a linear developmental relationship between clusters exists in males, as Tim4 and MHCII defined subsets were found to be replenished at similarly high rates. Hence, what dictates cluster identity in males remains unclear.

Given that the rate of replenishment from BM was markedly different between the sexes, this raised the possibility that transcriptional differences could reflect different ontogenies of male and female peritoneal macrophages. Indeed, a number of the genes we found to be expressed more highly by female peritoneal macrophages, including *Colec12*, *Cd163*, *Bmpr1a*, *Cdc42bpa, Timd4*, *Apoc1*, and members of the *Cd209* family have been reported to be expressed by embryonically, but not monocyte-derived macrophages in other tissues ^25, 55^. This could reflect an intrinsic property of their embryonic origin, or that such cells are likely to have resided in the tissue for a long period. However the fact that a proportion of BM-derived peritoneal macrophages can acquire the expression of at least some of these “embryonic” signature markers (e.g. Tim4, CD209b) in the setting of tissue-protected BM chimeras, suggests that this is more likely related to their time-of-residency rather than rigid differences related to origin. Consistent with this, co-expression of Tim4 and CD209b identifies the longest-lived macrophages in the peritoneal cavity irrespective of sex. The concept that macrophages require prolonged residence within the tissue to acquire their characteristic features is consistent with work from the Guilliams lab showing that acquisition of Tim4 expression by monocyte-derived cells that engraft in the liver following deletion of endogenous Kupffer cells increases with time ^25^. Notably, of the two populations of peritoneal macrophages that have been described in ascites fluid from patients with decompensated cirrhosis, the subset that aligns with mouse resident F4/80^hi^ macrophages exhibits significantly higher expression of *TIMD4*, *CD209*, *COLEC12*, CD163 and *APOC1* ^56^, suggesting these may represent phylogenetically conserved markers of long-lived macrophages.

We also identified features that mark newly differentiated CD102^+^ macrophages and are inversely related to time-of-residency, such as ApoE. This finding is consistent with repopulation studies showing that microglia of monocyte origin express higher levels of ApoE ^57–59^. While the association between *ApoE* and recent arrival seems to be at odds with the higher level of ApoE expression by female peritoneal macrophages, *ApoE* expression may also be controlled partly by estrogen, as the *Apoe* gene contains an estrogen response element, and its expression is reduced in inflammatory peritoneal macrophages by macrophage-specific deletion of the estrogen receptor alpha ^35^. Understanding how ApoE expression is regulated by tissue-residency and hormonal control may be important in many diseases in which is it is a known genetic risk factor and that exhibit strong sex-biases in risk, such as Alzheimer’s and cardiovascular disease ^60^, but also in healthy aging where *APOE* variants are among the strongest predictors of human longevity ^61^.

Importantly, we found the dimorphism in proliferation and CXCL13 expression by peritoneal macrophages to be regulated independently of macrophage replenishment kinetics, consistent with previous data showing that the proliferative capacity of macrophages is determined by signals in the local microenvironment rather than their origin ^25, 62^. Although these dimorphisms only developed following sexual maturation, it seems unlikely that estradiol levels are responsible for the lower proliferation of female peritoneal macrophages, as estradiol is reported to increase rather than inhibit proliferation of these cells ^36^. Furthermore, exogenous estradiol did not rescue the elevated turnover of female macrophages we found in ovariectomised mice. Similarly, exogenous estradiol does not influence CXCL13 expression by peritoneal macrophage ^36^, nor did it rescue the loss of B1 cells that occurred after ovariectomy (data not shown). While estradiol has been reported to upregulate the B1 cell regulators *Tnfsf13b*, *Tnfsf13,* and *Il10* ^15^by female peritoneal macrophages *in vitro*, none of these genes were differentially expressed by macrophages in our study. Moreover, genes highly expressed by male peritoneal macrophages, such as *Arg1* and *Chil3*, are increased by estradiol ^36^. Although expression of receptors for progesterone and androgens did not differ between the sexes, we cannot rule out a role for these steroids in generating sex dimorphisms. Thus, the exact local factor(s) driving the sex dimorphisms identified here remain to be elucidated. Interestingly, pathway analysis of our transcriptomic data identified ‘Interferon Gamma Response’ and ‘Interferon Alpha Response’ as gene sets enriched within female macrophages, and interferons are known to be hormonally regulated ^63^. Whether interferon receptor signalling plays a role in dimorphic characteristics of peritoneal macrophages is the focus of ongoing work.

The incidence and severity of sepsis and post-surgical infections are profoundly lower in women than men ^64^,but the mechanisms underlying these differences remain unclear. Our finding that female mice are more resistant to *S. pneumoniae* peritonitis is consistent with previous work on group B streptococcal peritonitis ^38^. However, while others attributed this to other elements of innate immune responses, such as neutrophil recruitment ^51^, our data suggest that the resistance of females is at least, in part, due to differences in resident peritoneal macrophages, such as elevated expression of CD209b. Dimorphic expression of CXCL13 may also contribute, as it plays a central role in recruiting B1 cells that produce the natural IgM ^2^ that protects against multiple forms of infectious peritonitis ^65, 66^. As activation of complement is essential for innate resistance to against *S.pneumoniae* }^65^ and both CD209b and natural IgM can activate the classical pathway of complement fixation during *S.pneumoniae* infection ^65–67^, peritoneal macrophages may play several, overlapping roles in protective immunity against this infection. Indeed *C1q*, *C3*, and *C4b*, as well as *Cfb,* which encodes factor B and is essential for the alternative pathway of complement fixation during *S.pneumoniae* infection, were also all expressed more highly by female peritoneal macrophages. We propose that this heightened barrier function in the female peritoneum may have evolved to mitigate the risk of sexually transmitted infection disseminating from the lower female reproductive tract ^68^ or to protect against puerperal peritonitis. Our findings also have wider implications for understanding peritoneal macrophage behaviour following a local mechanical or inflammatory insult, when tissue resident macrophages may be replaced by monocyte-derived cells that may require prolonged residence in the tissue before acquiring the full profile of resident macrophages with protective functions ^55, 69–71^. The potential risks in this process have been highlighted in the context of viral meningitis, where a failure of newly elicited macrophages to rapidly acquire CD209b expression led to impaired neutrophil recruitment to subsequent intra-cranial immune challenge ^72^, and could explain why animals exposed to sterile peritoneal inflammation are more susceptible to *S. pneumoniae* peritonitis for at least several months^73^.

Our studies highlight the importance of taking age and sex into account when understanding the peritoneal response to disease and implicate time-of-residency as an underlying determinant of resident macrophage function. Further work is needed to understand the molecular processes that underlie the requirement for time-of-residency on expression of these genes and to identify the local signals that govern this process. Beyond the cavity, our findings also have wider implications for the molecular mechanisms that drive dimorphic production of natural IgG by peritoneal B1 cells that provides women and infants with heightened resistance to blood-borne bacterial infections, particularly as these antibodies are lost in the absence of peritoneal macrophages ^15^.

## Materials and methods

### Animals and reagents

Wild type C57BL/6J CD45.2^+^, congenic CD45.1^+^CD45.2^+^ mice and *Ccr2^−/−^* mice were bred and maintained in specific pathogen-free facilities at the University of Edinburgh, UK. In some experiments, C57BL/6J (Crl) mice were purchased from Charles River, UK. *Itgax*^Cre 74^ (referred here to CD11c^Cre^) mice were crossed with *Rosa26*^LSL-YFP^ mice (a gift from Dr. Megan Mcleod, University of Glasgow, UK) and maintained at the University of Glasgow. For *Cdh5*^Cre-ERT2^ fate mapping, WT females aged 6-10 weeks were subjected to timed matings with *Cdh5*^CreERT2+/−^ or *Cdh5*^CreERT2+/+^ *Rosa26*^tdT/tdT^ males. Successful mating was judged by the presence of vaginal plugs the morning after, which was considered 0.5days post conception. For induction of reporter recombination in the offspring, a single dose of 4-hydroxytamoxifen (4OHT; 1.2mg) was delivered by i.p. injections to pregnant females at E7.5. To counteract adverse effects of 4OHT on pregnancy, 4OHT was supplemented with progesterone (0.6mg). In cases when females could not give birth naturally, pups were delivered by C-section and cross-fostered with lactating CD1 females. All experimental mice were age and sex matched. To perform estrous staging, vaginal lavage was performed and cellular content examined following haematoxylin and eosin staining, as previously described ^75^. For high fat diet (HFD) experiments, tissue protected BM chimeric mice were placed on HFD (58 kcal% fat and sucrose, Research Diet, D1233li) for 8 weeks starting 4 weeks post reconstitution. Experiments performed at UK establishments were permitted under license by the UK Home Office and were approved by the University of Edinburgh Animal Welfare and Ethical Review Body or the University of Glasgow Local Ethical Review Panel.

### Surgery

Ovariectomy/oophorectomy was performed on 6-week-old wild type (C57BL/6J) or tissue protected BM chimeras 8 weeks post-reconstitution (16 weeks of age). Briefly, dorsal unilateral or bilateral ovariectomy (OVX) was performed and mice allowed to recover for up to 8 weeks. Sham surgery was performed to control for the effects of surgery on the peritoneal environment. This involved identical surgery except for the excision of the ovary/ovaries. Surgery was performed under isoflurane anaesthesia followed by a postoperative analgesic, buprenorphine (0.1 mg/kg), for pain management. In some experiments following 7 days of recovery, mice received exogenous estradiol in the form of E2 valerate (E2; 0.01 mg/kg) s.c. thrice weekly for 3 weeks.

### Tissue-protected BM chimeric mice

Anaesthetised 6-12 week old C57BL/6J CD45.1^+^CD45.2^+^ animals were exposed to a single dose of 9.5 Gy *γ*-irradiation, with all but the head and upper thorax of the animals being protected by a 2 inch lead shield. Animals were subsequently given 2-5×10^6^ BM cells from sex matched or mismatched congenic CD45.2^+^ C57BL/6J animals by i.v. injection. Unless specified, mice were left for a period of at least 8 weeks before analysis of chimerism in the tissue compartments.

### BrdU injection

For labelling of proliferating cells, mice were injected s.c. with 100μl of 10mg/ml BrdU (Sigma) in Dulbecco’s PBS 2hr before culling.

### Preparation of single cell suspensions

Mice were sacrificed by CO_2_ inhalation or by terminal anaesthesia followed by exsanguination. Peritoneal and pleural cavities were lavaged with RPMI containing 2mM EDTA and 10mM HEPES (both Invitrogen) as described previously ^76^. Any samples with excessive erythrocyte contamination were excluded from analysis. Omental tissue was excised, chopped finely and digested in 0.5ml pre-warmed collagenase D (1mg/ml; Roche) in RPMI 1640 media supplemented with 2% FCS for 15 minutes in a shaking incubator at 37°C. Following disaggregation with a P1000 Gilson, omental tissue was digested for a further 20mins before being placed on ice. 2.5µl of 0.5M EDTA was added to each sample to inhibit enzymatic activity. Cell suspensions were passed through an 100μm filter and centrifuged at 1700rpm for 10mins. The resulting cell suspension was subsequently passed through a 40μm strainer prior to cell counting. All cells were maintained on ice until further use. Cellular content of the preparations was assessed by cell counting using a Casey TT counter (Roche) in combination with multi-colour flow-cytometry.

### Flow cytometry

Equal numbers of cells were blocked with 0.025 μg anti-CD16/32 (2.4G2; Biolegend) and 1:20 heat-inactivated mouse serum (Invitrogen), and then stained with a combination of the antibodies detailed in Supplementary Table 2. Where appropriate, cells were subsequently stained with streptavidin-conjugated fluorochromes. Dead cells were excluded using DAPI, 7-AAD or Zombie Aqua fixable viability dye (Biolegend). Fluorescence-minus-one controls confirmed gating strategies, while discrete populations within lineage^+^ cells were confirmed by omission of the corresponding population-specific antibody. Erythrocytes in blood samples were lysed using 1x RBC Lysis buffer (Biolegend), as per the manufacturer’s guidelines. For intracellular staining, cells were subsequently fixed and permeabilized using FoxP3/Transcription Factor Staining Buffer Set (eBioscience), and intracellular staining performed using antibodies detailed in Supplementary Table 2. For the detection of BrdU, cells were fixed as above and incubated with 3μg DNaseI (Sigma) for 30-60mins, before being washed in PermWash (eBioscience) and then incubated with anti-BrdU antibody for 30mins at RT. Samples were acquired using a FACS LSRFortessa or AriaII using FACSDiva software (BD) and analyzed with FlowJo software (version 9 or 10; Tree Star). Analysis was performed on single live cells determined using forward scatter height (FCS-H) versus area (FSC-A) and negativity for viability dyes. For analysis of macrophage proliferation, Ki67 expression was used to determine the frequency of all CD102^+^/F4/80^hi^ cells in cycle, whereas a 2h BrdU pulse before necropsy combined with Ki67 expression was used to identify cells in S phase, as described previously ^77^. mRNA was detected by flow cytometry using PrimeFlow technology (ThermoFisher) using probes against ApoE (probe type 10; AF568), ApoC1 (probe type 4; AF488) and CXCL13 (probe type 6; AF750) according to the manufacturer’s guidelines.

### Transcriptional Analysis

#### Bulk RNAseq

CD102^+^F4/80^hi^ cells were FACS-purified from the peritoneal and pleural cavities of unmanipulated male and female mice. For each population, 25,000 cells were sorted into 500μl RLT buffer (Qiagen) and snap frozen on dry ice. RNA was isolated using the RNeasy Plus Micro kit (Qiagen), at which point triplicates of 25,000 cells for each population were pooled. 10 ng of total RNA were amplified and converted to cDNA using Ovation RNASeq System V2 (Nugen). Sequencing was performed by Edinburgh Genomics using the Illumina HiSeq 4000 system (75PE). Raw map reads were processed with the R package DESeq2 ^78^ to generate the differentially expressed genes, and the normalized count reads to generate and visualize on heat maps generated by the R package pheatmap. Samples with >5% of reads mapped to ribosomal RNA were removed from analysis. DEG were determined using at least a 1.5-fold difference and adjusted *p* < 0.01, for each of the six pairwise comparisons. Pathway enrichment analysis was performed using the GSEA online database and the R package gskb (Gene Set data for pathway analysis in mouse) which makes predictions between each of the six pairwise comparisons, incorporating in the analysis the statistically significant differences in gene expression. The R package gskb was used to determine the chromosomal location of each of the genes and transcription factors. All R code is available upon request.

#### Single-cell RNAseq

10K cells for male and female sorted cells were loaded in Chromium 10x in parallel. Libraries were prepared as per manufacturer’s protocol and sequenced on Illumina Novaseq S1. Initial processing was done using Cellranger (v2.1.1) mkfastq and count (aligned to mouse assembly mm10). ***Preparation of analysis ready data:*** For each dataset (filtered data from Cell Ranger pipeline), we filtered out potentially low quality cells using dataset-specifc thresholds based on the trend of the number of genes per cell versus number of housekeeping genes per cell and number of genes per cell versus percentage of mitochondrial genes per cell curves as follows. More specifically, for the female data, we retained 4341 cells that have between 300 and 5000 genes, at least 65 housekeeping genes and percentage of mitochondrial genes over the total number of expressed genes below 2%. For the male data, we retained 2564 cells that have between 300 and 6000 genes, at least 70 housekeeping genes and percentage of mitochondrial genes over the total number of expressed genes below 2%. Finally, we filtered out genes that were expressed in less than 1% of the cells from each dataset. ***Clustering analysis of the data:*** Clustering and data merging using CCA was done using Seurat (v3.1.0). We used default parameters and 20 principal components for aligning and clustering the data. We next removed a very small cluster that lay far from all other clusters on the UMAP projection, indicating it could be either contamination or doublets and constructed a phylogenetic tree of the remaining clusters to understand the distances and relationship between them. Clusters that were closely grouped together and did not show unique markers, were merged together. The final result consists of 6890 cells grouped into 6 clusters. ***Identification of differentially expressed genes in CCA aligned clusters:*** We used MAST (v1.10) as implemented in the Seurat package and with default parameters to identify differentially upregulated genes between the identified clusters. To overcome the bias of batch effect, we found differentially upregulated genes within each dataset separately and retained the intersection of markers (conserved markers). ***Identification of differentially expressed genes between female and male cells:*** We used Student’s t-test as implemented in the Seurat package between equivalent female and male cells to identify differentially upregulated genes between male and female cells. We only retained genes with adjusted p-value based on Bonferroni correction below 0.05. Genes that were identified as differentially expressed for more than four out of the six clusters were selected as global DE genes and were removed from the cluster-specific differences. ***Pseudotemoral orderning:*** We used monocle (v2.12.0) with default parameters to build pseudotemporal trajectories of the female and the male data separately. ***Functional annotation of genesets***: We used DAVID to obtain enriched Gene Ontology Biological Process (GO-BP) and KEGG pathway terms for each extracted geneset. Given a list of genes upregulated in a set of cells (e.g. cluster) as target and a list of all genes observed in the respective dataset as background, we downloaded a list of GO-BP terms (GOTERM_BP_FAT) and a list of KEGG pathway terms that were significantly enriched in the target list as a functional annotation chart. We kept only terms with Benjamini adjusted p-values less than 0.05 and sorted them by decreasing Fold Enrichment.

### S. pneumoniae peritonitis

*S. pneumoniae* were cultured overnight on blood agar plates (5% CO_2_, 95% air, 37 C), inoculated into Brain Heart Infusion broth, cultured for 3 h, washed, and resuspended at 10^4^ CFU/ ml (estimated by OD_595_) in sterile PBS. Their concentration was verified by serial dilution and culture on blood agar plates. Groups of male and female, age-matched C57Bl/6 mice (8–14 wk of age) were inoculated intraperitoneally with 100 μl of PBS containing 10^3^ CFU *S. pneumoniae* (capsular type 2 strain D39). Mice were culled 20 h later and peritoneal lavage performed using sterile PBS. 100 μl of lavage fluid was cultured for bacterial growth for 24 h. The remaining lavage fluid was centrifuged at 400g for 5 mins and the resulting cells counted and prepared for flow cytometric analysis.

### Statistics

Statistics were performed using Prism 7 (GraphPad Software). The statistical test used in each experiment is detailed in the relevant figure legend.

## Accession codes

### Data availability

Data that support the findings of this study are available from the corresponding authors upon reasonable request.

## Acknowledgements

Flow cytometry data were generated with support from the QMRI Flow Cytometry and Cell Sorting Facility, University of Edinburgh. mRNA sequencing was performed by Edinburgh Genomics, The University of Edinburgh. Edinburgh Genomics is partly supported through core grants from NERC (R8/H10/56), MRC (MR/K001744/1) and BBSRC (BB/J004243/1). Servier Medical Art was used for the generation of some of the graphics.

This work was funded by the Medical Research Council UK (MR/L008076/1 to S.J.J), with additional support from the Wellcome Trust (IS3-R34 to S.J.J; PhD studentship 203909/Z/16/A to P.L.) and a Sir Henry Dale Fellowship jointly funded by the Wellcome Trust and the Royal Society (Grant Number 206234/Z/17/Z to C.C.B).

## Author Contributions

C.C.B. conceived and performed most of the experiments, analysed and interpreted the data, wrote the manuscript and provided funding. D.A.G. designed and performed experiments and edited the manuscript. N.S. performed transcriptomic analysis (population level RNAseq) and figure generation. K.B. performed single cell RNAseq analysis and figure generation. P.L. performed experiments for generation of scRNAseq data. C.D. provided technical assistance for the design and execution of infection experiments. R.G. performed *Cdh5*-fate mapping experiments. M.M-P. helped with the design and execution of high fat diet experiments. C.B. helped with the design and execution of high fat diet experiments. M.B. provided access to *Cdh5* fate-mapper mice. D.D. helped with the design and execution of infection experiments. P.T.K.S. provided advice for the design and interpretation of experiments and edited the manuscript. N.B. performed the scRNAseq analysis, provided advice on the interpretation of these data, edited the manuscript and provided funding. S.J.J. conceived and performed experiments, analysed and interpreted the data, wrote the manuscript, obtained funding and supervised the project.

---

**Supplementary Figure 1**

(**a**) Normalized non-host chimerism of peritoneal F4/80^hi^CD102^+^ macrophages and indicated leukocytes from sex matched tissue-protected BM chimeric mice 8-12 weeks post-reconstitution. Data are normalised to the non-host chimerism of Ly6C^hi^ monocytes. Symbols represent individual animals and horizontal bars represent the mean. Data represent 10 mice per group pooled from two independent experiments. ****P<0.0001. Student’s *t* test with Holm-Sidak correction. (**b**) Gating strategy for the identification of omental macrophage subsets. (**c**) Expression of MHCII by CD102-defined macrophage subsets and CD11b^+^ DC from omental digests. (**d**) Normalized non-host chimerism of peritoneal and omental CD102^+^ macrophages from sex matched and sex mismatched tissue-protected BM chimeric mice 8-12 weeks post-reconstitution. Data are normalised to the non-host chimerism of Ly6C^hi^ monocytes. Symbols represent individual animals and horizontal bars represent the mean. Data represent 9 (sex mismatched) or 10 (sex matched) mice per group pooled from two independent experiments. **P<0.01. Student’s *t* test with Holm-Sidak correction.

**Supplementary Figure 2**

(**a, b**) Frequency of F4/80^hi^CD102^+^ macrophages (**a**) and CD102^−^MHCII^+^ cells (**b**) of total CD45^+^ cells obtained from the peritoneal cavity of tissue-protected BM chimeric mice that had received surgery 8 weeks earlier. (**c-f**) Frequency (*left panels*) and absolute number of eosinophils (**c**), B1 cells (**d**), B2 cells (**e**) and T cells (**f**) from the peritoneal cavity of tissue-protected BM chimeric mice that had received surgery 8 weeks earlier. Symbols represent individual animals and horizontal bars represent the mean. Data represent 9 (sham) or 10 (control, unilateral, bilateral) mice per group pooled from two independent experiments.

**Supplementary Figure 3**

(**a**) Experimental scheme for the generation of sex matched, tissue-protected bone marrow (BM) chimeric mice and high-fat diet regimen. (**b**) Mass of abdominal adipose tissue in male and female mice fed control or high fat diet for 4 weeks and rested for 8 weeks. (**c**) Bodyweight of male and female mice fed control or high fat diet for 4 weeks and rested for 8 weeks (8) paired to starting bodyweight (0). (**d**) Normalized non-host chimerism of peritoneal F4/80^hi^CD102^+^ macrophages from mice in **a.** Symbols represent individual animals and horizontal bars represent the mean. Data represent 5 (**b**) mice from one experiment or 10 (**c, d**) mice per group pooled from two independent experiments.

**Supplementary Figure 4**

(**a**) Gating strategy for the identification and FACS-purification of CD102^+^ macrophages for population-level RNAseq.

(**b**) Expression of F4/80 by peritoneal CD102-defined macrophages.

(**c**) Representative post-sort purity of CD102-defined macrophages.

**Supplementary Figure 5**

(**a**) Expression of indicated TLRs and TLR adaptor molecules by male and female peritoneal macrophages in the population-level RNAseq data set. (**b**) Expression of selected genes by male and female peritoneal macrophages that were identified by Villa et al. (Cell Reports) to be expressed in a sexually dimorphic fashion by microglia.

**Supplementary Figure 6**

(**a**) Representative expression of CD102 and CD209b by peritoneal all CD45^+^ leukocytes obtained from age-matched male and female C57BL/6 mice. (**b**) Representative expression of Tim4 and CD209b by peritoneal CD102^+^ macrophages obtained from age-matched male and female C57BL/6 mice housed at the indicated institutions. (**c**) Representative expression of Tim4 and CD209b by peritoneal CD102^+^ macrophages obtained from age-matched male and female *Rag1^−/−^* mice (*left*) and frequency of CD209b^+^Tim4^+^ macrophages, the mean fluorescence intensity (MFI) of CD209b and frequency of Tim4*^−^* macrophages in each sex. Data are from one experiment with 4 (female) or 6 (male) mice per group. ** P<0.01, ***P<0.001, ****P<0.0001. Student’s *t* test. (**d**) Representative expression of CD209b, Tim4 and Ki67 by peritoneal CD102^+^ macrophages obtained from age-matched male and female BALB/c mice (*left*) and a summary of replicate data showing the frequency of CD209b^+^, Tim4^+^ and Ki67^+^ cells as well as MFI of expression. Data are from one experiment with 5 (female) or 6 (male) mice per group. *P<0.05, ** P<0.01, ***P<0.001, ****P<0.0001. Student’s *t* test.

**Supplementary Figure 7**

(**a**) Absolute number of FRβ^+^ CD102^+^ macrophages obtained from the peritoneal cavity of unmanipulated 22-28 week old *Ccr2*^+/+^ or *Ccr2*^−/−^ mice. Symbols represent individual animals and horizontal bars represent the mean. Data are from one experiment with 2 (*Ccr2*^+/+^ males), 4 (*Ccr2*^+/+^ females) or 5 (*Ccr2*^−/−^) mice per group. (**b**) Absolute number of B1 cells obtained from the peritoneal cavity of unmanipulated male and female mice at 6-8 weeks and 14-16 weeks. Symbols represent individual animals and horizontal bars represent the mean. Data are pooled from four experiments with 9 (females, both time points), 8 (male, 6-8 weeks) or 13 (male, 14-16 weeks) mice per group. Data for the 14-16 week time point taken from data in Figure 6g. ****P<0.0001. Two-way ANOVA.

