## Supplemental Figures for "Origin and microenvironment contribute to the sexually dimorphic phenotype and function of peritoneal macrophages"

### Supplementary Figure 1

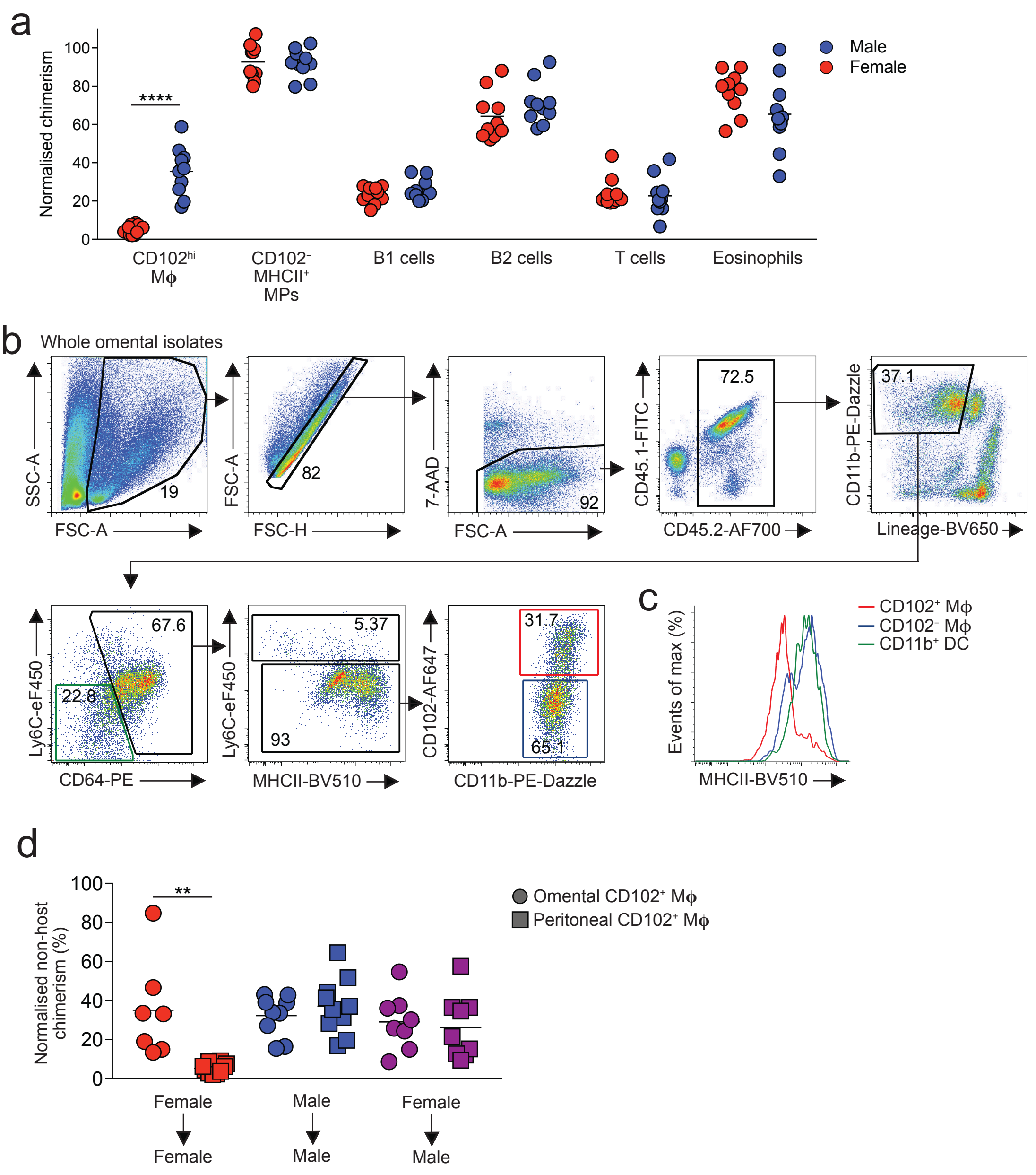

### Supplementary Figure 2

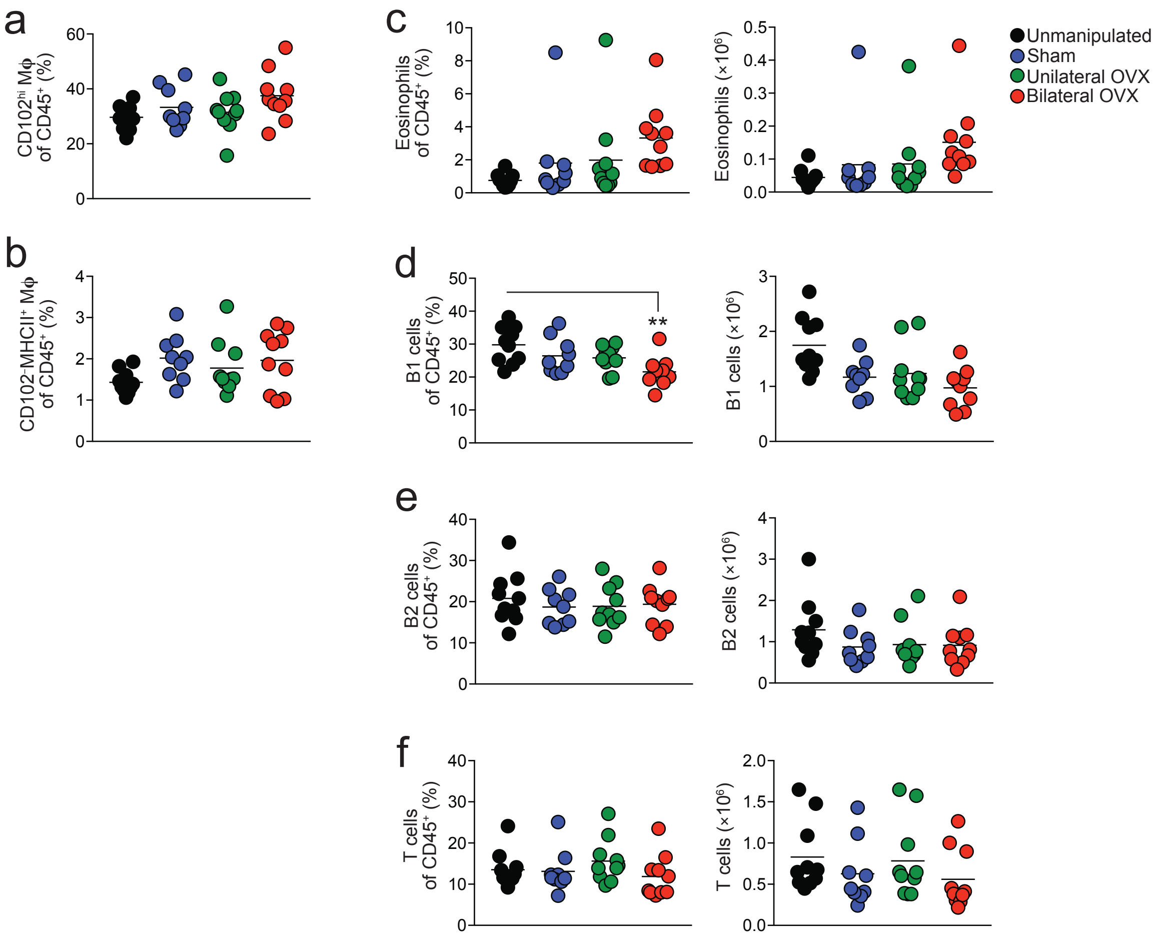

### Supplementary Figure 3

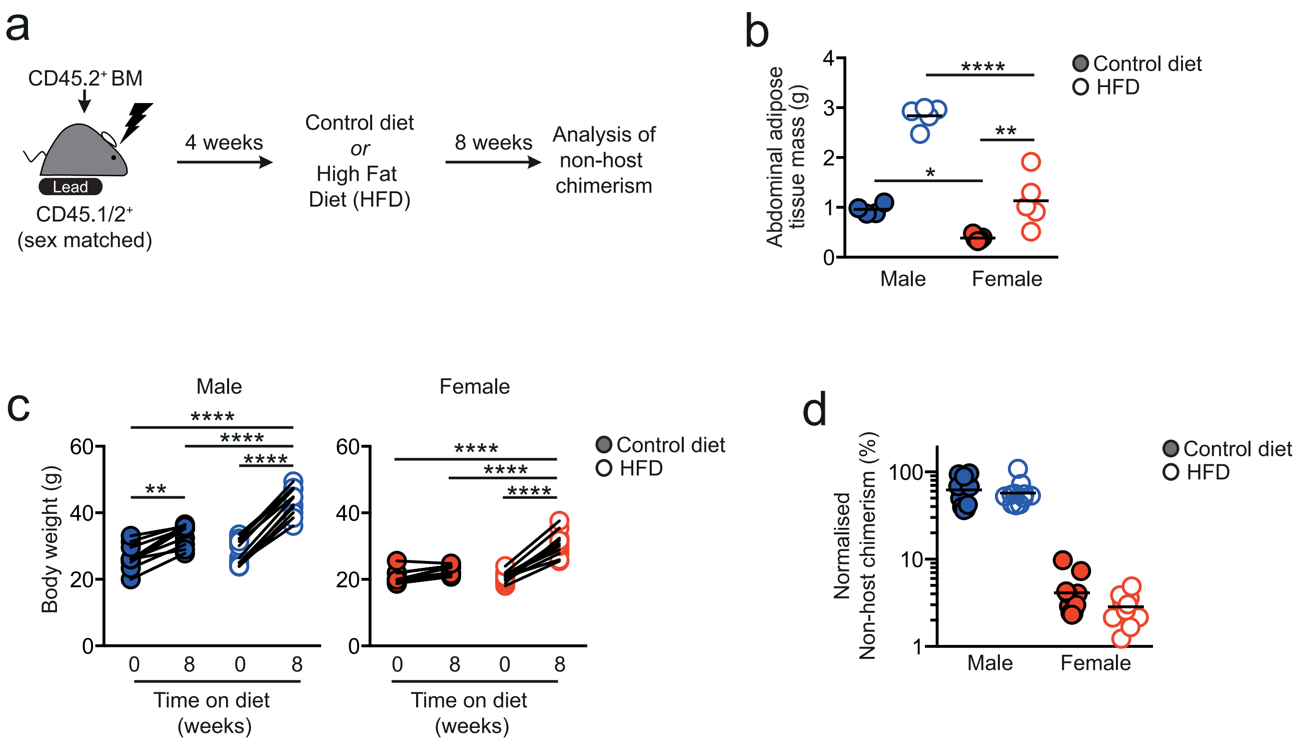

### Supplementary Figure 4

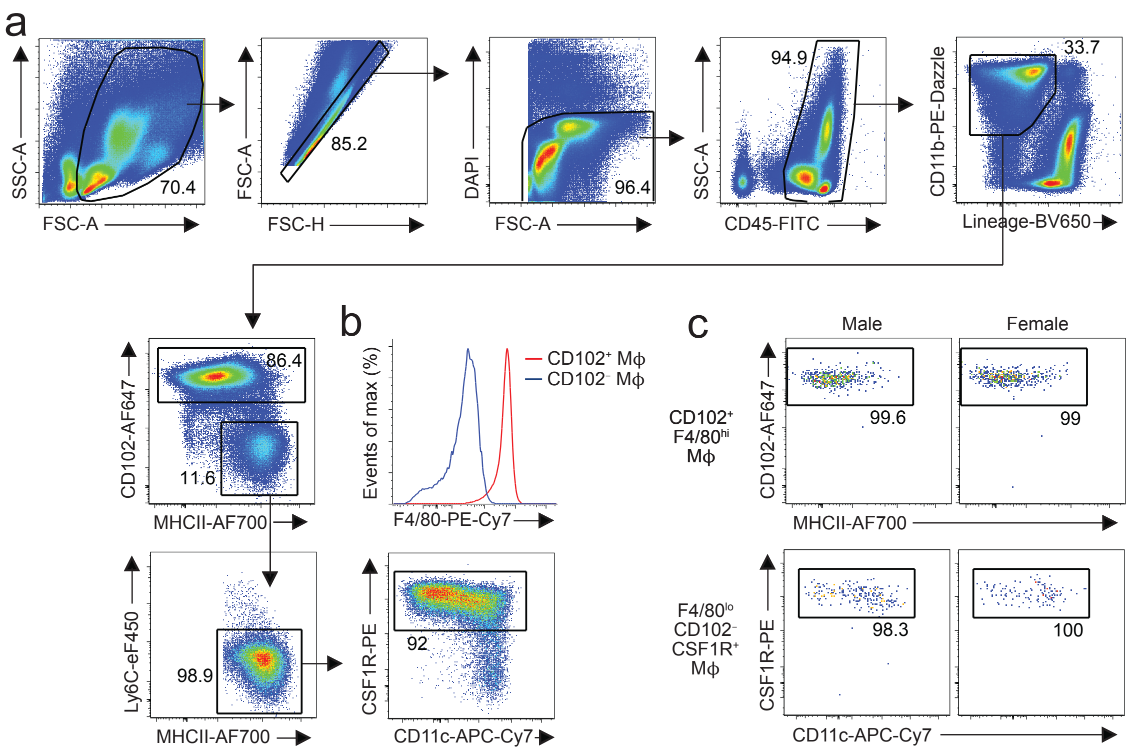

### Supplementary Figure 5

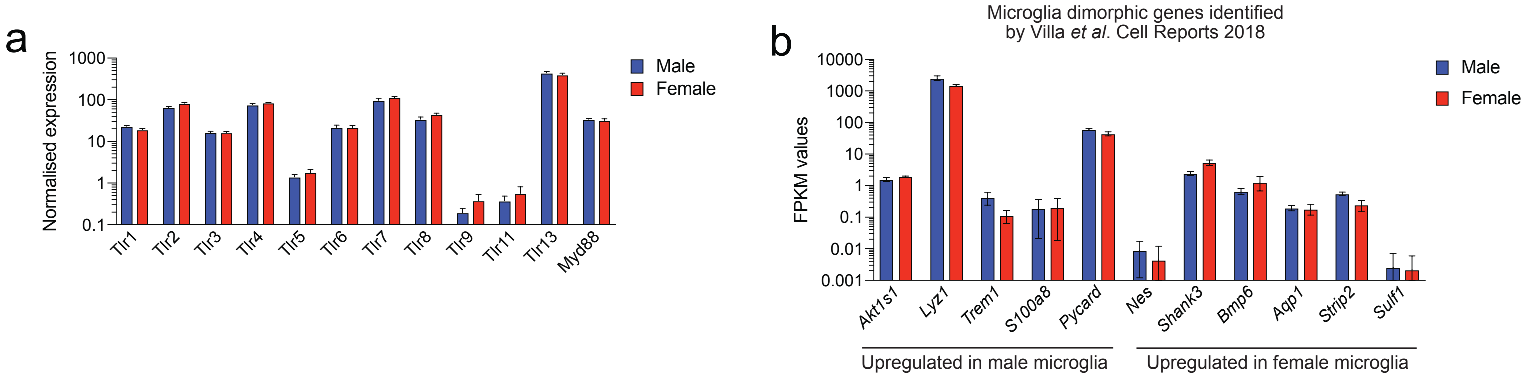

### Supplementary Figure 6

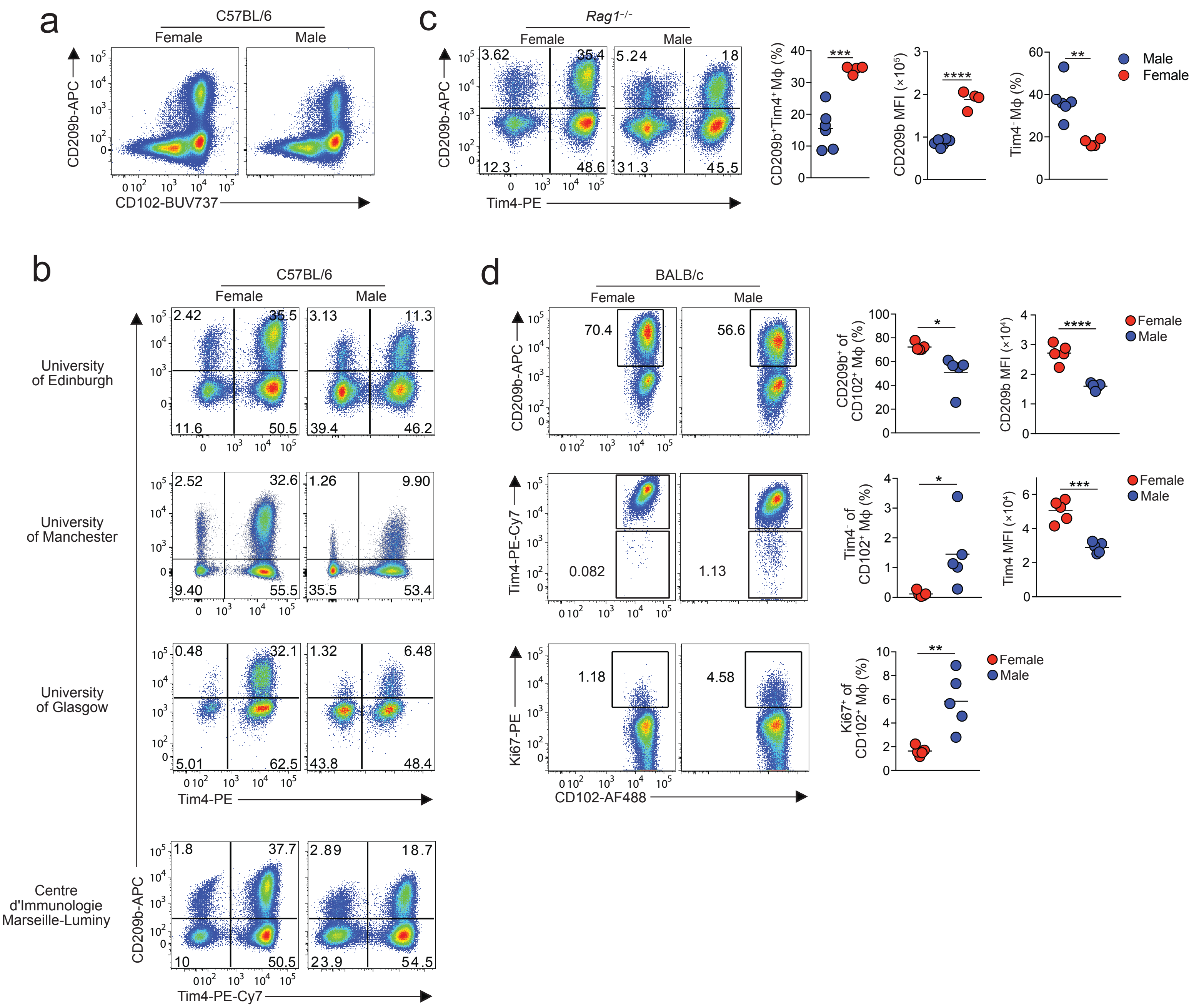

### Supplementary Figure 7

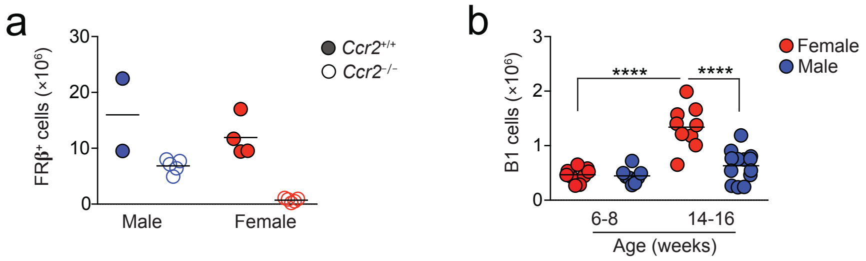
