## Supplemental Table 6 for "Origin and microenvironment contribute to the sexually dimorphic phenotype and function of peritoneal macrophages"

**Supplementary Table 2**

| **Antibody** | **Clone** | **Source** | **Fluorochrome** | **Catalogue #** |
| --- | --- | --- | --- | --- |
| **Mouse** |  |  |  |  |
| BrdU | Bu20a | Biolegend | PE | 339812 |
| CD3 | 17A2 | Biolegend | Biotin | 100244 |
| CD11b | M1/70 | Biolegend | PE-Dazzle | 101256 |
|  |  |  | APC/Fire 750 | 101256 |
| CD11c | N418 | Biolegend | APC-Cy7 | 117324 |
| CD14 | Sa2-8 | eBioscience/ThermoFisher | APC | 12-0141-82 |
| CD16/32 | 2.4G2 | Biolegend | Purified | 101320 |
| CD19 | 6D5 | Biolegend | Biotin | 115504 |
| CD45 | 30-F11 | Biolegend | FITC | 103108 |
| CD45.1 | A20 | Biolegend | FITC | 110706 |
| CD45.2 | 104 | Biolegend | AF700 | 109822 |
| CD62L | MEL-14 | Biolegend | PE | 104408 |
| CD63 | NVG-2 | Biolegend | PE | 143904 |
| CD64 | X54-5/7 | Biolegend | PE | 139304 |
| CD102 | 3C4 | Biolegend | AF488 | 106065 |
|  |  |  | AF647 | 105612 |
|  |  |  | Biotin | 105604 |
| CD115 | AFS98 | Biolegend | PE | 135505 |
|  |  |  | APC | 135510 |
| CD209b | 22D1 | eBioscience/ThermoFisher | APC | 17-2093-82 |
| GATA6 | D61E4 | Cell Signalling Technologies | Purified | 5851S |
| F4/80 | BM8 | Biolegend | PE-Cy7 | 123114 |
|  |  |  | APC | 123116 |
|  |  |  | BV785 | 123141 |
| Ki67 | B56 | BD Biosciences | AF488 | 558616 |
|  | REA183 | Miltenyi Biotec | FITC | 130-117-803 |
| Ly6C | HK1.4 | eBioscience/ThermoFisher | eFluor450 | 48-5932-82 |
| Ly6G | 1A8 | Biolegend | Biotin | 127604 |
| MHC II (IA-IE) | M5/114.15.2 | Biolegend | AF700 | 107622 |
|  |  |  | BV510 | 107636 |
| RELMα | Polyclonal | Peprotech | Purified | 500-P214-100 |
| SiglecF | E50-2440 | BD Biosciences | PE-CF594 | 562757 |
| SiglecF | ES22-10D8 | Miltenyi Biotec | Biotin | 130-101-861 |
| Tim4 | RMT4-54 | Biolegend | PE | 130006 |
|  |  |  | PE-Cy7 | 130010 |
|  |  |  | AF647 | 130008 |
| VSIG | NLA14 | eBioscience/ThermoFisher | PE-Cy7 | 25-5752-82 |
| Streptavidin |  | Biolegend | BV650 | 405232 |
| Zenon anti-rabbit reagent | | Molecular Probes | AF488 | Z25302 |
|  | |  | PE | Z25355 |
